# STING–IFN–CH25H lipid axis links innate immune activation to tau pathology

**DOI:** 10.64898/2025.12.16.694702

**Authors:** Hao Chen, Li Fan, Man Ying Wong, Yohannes A. Ambaw, Jingjie Zhu, Kendra Norman, Pearly Ye, Nessa Foxe, Daphne Zhu, Yuansong Wan, Mahalashmi Srinivasan, Sue-Ann Mok, Laura Beth J McIntire, Tobias C. Walther, Robert V. Farese, Li Gan, Wenjie Luo

## Abstract

Genetic risk for Alzheimer’s disease is strongly enriched in pathways governing microglial activation and cholesterol metabolism, yet how these processes converge to drive neurodegeneration remains unclear. Here, we identify the oxysterol 25-hydroxycholesterol (25- HC) as a pathogenic lipid downstream of the cGAS-STING–IFN pathway. In models of tau pathology, interferon signaling induces Cholesterol 25-hydroxylase (CH25H) expression in microglia. Ch25h deletion in female P301S mice suppressed tau aggregation, preserved synapses, prevented brain atrophy, and rescued memory. Mechanistically, loss of CH25H disrupted STING trafficking, attenuated IFN activation, dampened self-perpetuating microglial inflammation. Strikingly, 25-HC directly accelerated tau propagation in human iPSC derived neurons. It also disrupted lysosomal and mitochondrial lipid composition, driving cholesteryl ester accumulation and promoting apoptosis under tau-induced stress. These findings define a STING–IFN–CH25H lipid axis that bridges innate immune activation to tau pathology and toxicity, offering a tractable therapeutic pathway for inflammation-driven neurodegenerative conditions.

## Introduction

Alzheimer’s disease (AD) is characterized by the accumulation of amyloid-β plaques and tau neurofibrillary tangles, accompanied by profound remodeling of the brain’s innate immune landscape^1–4^. Among the most dynamic responders are microglia, which undergo extensive transcriptomic reprogramming into disease-associated microglia (DAM)^5^ or microglia with neurodegenerative phenotype (MGnD)^6,7^. Microglia in diseases display broad alterations in immune signaling, lipid metabolism, and phagocytic capacity, underscoring the intimate coupling between lipid homeostasis and immune activation in the degenerating brain^7–14^. Increasing evidence suggests that this relationship is bidirectional: lipid species modulate immune responses, while immune activation reshapes lipid metabolism to sustain or amplify inflammation^10,15–18^. Little is known about the molecular drivers of the bidirectional relationships, and how they influence neuronal vulnerability.

Activated microglia release a wide spectrum of effector molecules that influence neurons and other glial cells^15,16,19–21^. While cytokines such as IL-1β and type I interferons have been extensively studied, non-cytokine mediators, including lipids, metabolites, and small signaling molecules, are emerging as equally important yet poorly defined contributors to neurodegeneration. *CH25H* is a microglial-enriched gene upregulated in DAM in AD and related tauopathies^5,7,22–26^. This enzyme catalyzes the oxidation of cholesterol to produce 25-hydroxycholesterol (25-HC), an oxysterol known to regulate cholesterol turnover and modulate both innate and adaptive immune signaling in a context-dependent manner^27–32^. Expression of *CH25H* is strongly induced by type I interferons (IFN-I) and other proinflammatory cues, including LPS and demyelination ^14,29,33–37^, positioning it as a potential downstream effector of IFN signaling. Through this IFN-driven induction, CH25H acts as an effector of the antiviral response, restricting viral replication while simultaneously shaping inflammatory tone^35–40^. Emerging studies suggest that IFN-driven CH25H upregulation in microglia may represent a mechanism of lipid-mediated immune modulation^22,38,41^. Elevated 25-HC can influence microglial phenotype and inflammatory tone, potentially amplifying or constraining neuroinflammatory responses depending on disease stage and local context^22,39,42–46^. In AD, chronic IFN signaling and sustained CH25H activation could thus contribute to maladaptive microglial activation and neurodegeneration.

Our prior work demonstrated that 25-HC augments IL-1β release by activating inflammasomes in microglia ^22^, suggesting a proinflammatory role in the aging and diseased brain. Consistent with its proinflammatory properties, recent studies have linked 25-HC to enhanced tau pathology^47^ or amyloid pathology^41^ in vivo. Moreover, 25-HC may act as a key mediator of microglial disruption of hippocampal plasticity^46,48^, implicating this oxysterol in the neurodegenerative cascade. However, the mechanistic pathways through which 25-HC contributes to microglial activation, how these interactions shape neuronal lipid composition and tau propagation, have not been resolved.

Here, we demonstrate that *Ch25h* deletion in female tauopathy mice mitigates tau-induced neurodegeneration by reducing neuronal tau pathology and brain atrophy, while shifting brain transcriptome from proinflammatory toward homeostatic states. Our results reveal a self-reinforcing inflammatory loop in microglia with Ch25h required for STING activation. We further identify a direct neuron-specific action of 25-HC in facilitating tau propagation and promoting neuronal apoptosis, accompanied by reprogramming of neuronal lipids, including elevating cholesterol esters, reducing lysosomal bis(monoacylglycerol) phosphate (BMP) and mitochondrial cardiolipin (CL). These findings uncover the STING–CH25H–25-HC axis as a pivotal mechanism in tau-driven neurodegeneration and identify it as a compelling therapeutic avenue for tauopathy.

## Results

### Ch25h deficiency ameliorates tau pathology and neurodegeneration in female tauopathy mice

To investigate the role of CH25H in tau pathogenesis, we crossed *Ch25h*^-/-^ mice with the human tau transgenic mice expressing human 1N4R Tau with the *P301S* mutation (PS19 mice), a tauopathy model developing age-dependent tau pathology and severe neurodegeneration ^49^. At 9 months of age, we found that female *Ch25h^-/-^ Tau^+^* mice exhibited significant less tau pathology in the hippocampal CA1 and Dental Gyrus (DG) regions compared to *Ch25h^+/+^ Tau^+^* littermates, as detected by immunostaining using the conformational-specific tau antibody MC-1 (**Fig. 1A-C, supplementary dataset-1**). Deletion of *Ch25h* also markedly attenuated tau-induced lateral ventricular enlargement, as assessed by Sudan black staining (**Fig. 1D-E, supplementary dataset-1**). Consistent with these findings, plasma levels of neurofilament light chain (Nfl), a well-established biomarker of neuronal damage, were significantly lower in *Ch25h^-/-^ Tau^+^* mice than in controls (**Fig. 1F, supplementary dataset-1**). In contrast, *Ch25h* deletion did not alter ventricular volume or plasma Nfl levels in male tauopathy mice (**Supplementary Fig. 1A-B, supplementary dataset-1**), suggesting a possible sex specific effect. Based on these observations, subsequent analyses focused primarily on female mice.

**Figure 1:**
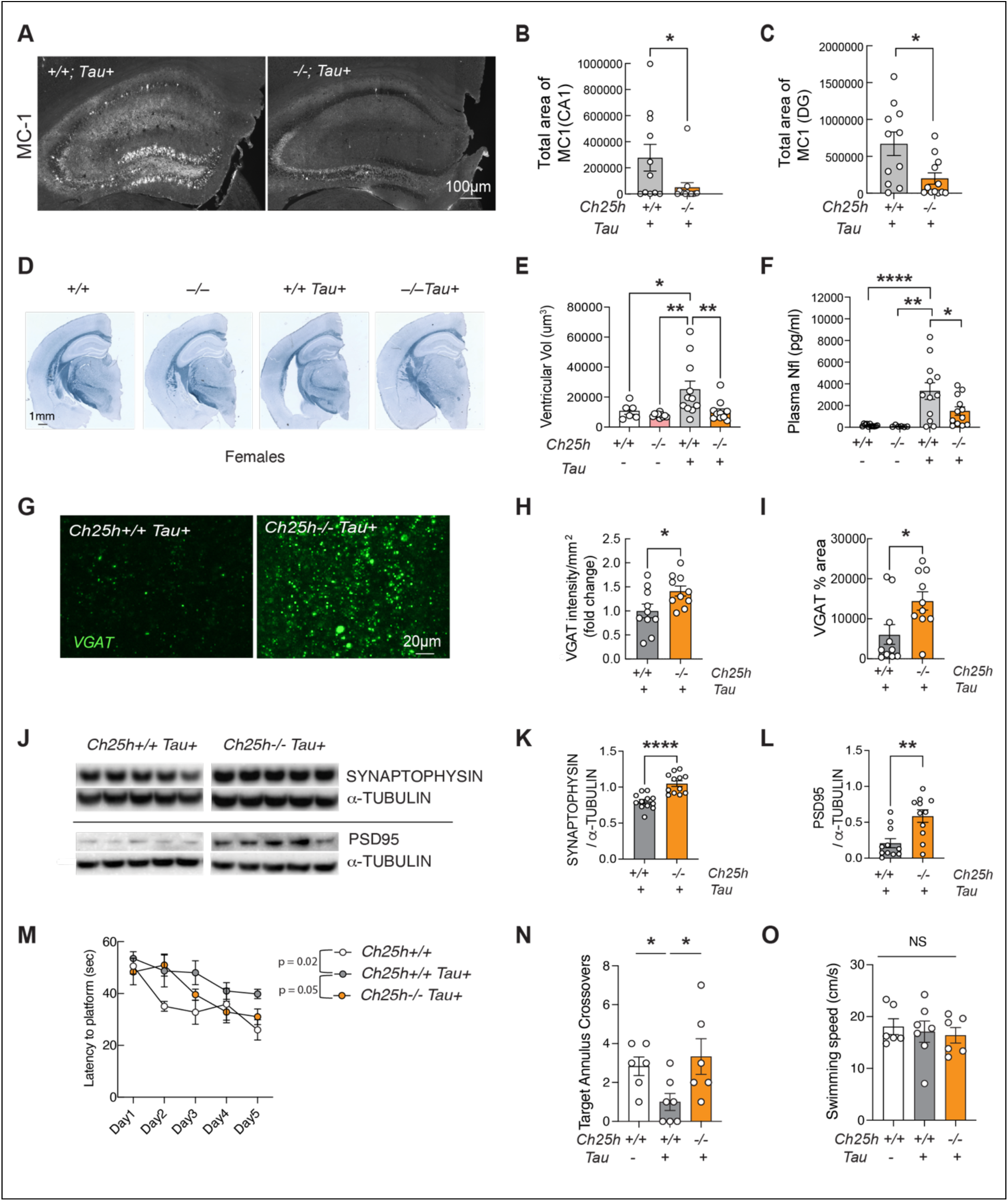
C*h*25h deficiency mitigates tau pathology, neurodegeneration and cognitive deficit in female tauopathy mice. A) Representative images of MC1 staining in hippocampal area of female *Ch25h+/+ Tau+* and *Ch25h-/-Tau+* mice. scale bar =100µm B-C) Quantification of MC1+ area in CA1 region (B) and in dentate gyrus (DG) (C). Unpaired student’s t-test. *p<0.05. n= 11 mice for *Ch25h+/+ Tau+*, n = 12 mice for *Ch25h-/-Tau+*. D) Representative images of brains from 9- to 10-month-old female mice stained with Sudan black. E) Quantification of ventricle volumes across four genotypes. Two-Way ANOVA followed by post-hoc Tukey test. *p<0.05, **p<0.01. n = 6 mice for *Ch25h+/+*, n = 9 mice for *Ch25h-/-,* n = 11 mice for *Ch25h+/+ Tau+*, n = 10 mice for *Ch25h-/-Tau+*. F) Plasma neurofilament Light (Nfl) levels across four genotypes. Two-Way ANOVA followed by posthoc Tukey test. *p<0.05, **p<0.01, ****p<0.0001. n = 6 for *Ch25h+/+* mice, n = 9 mice for *Ch25h-/-,* n = 11 mice for *Ch25h+/+ Tau+*, n = 12 mice for *Ch25h-/-Tau+*. G) Representative images of VGAT staining in CA1 region of female *Ch25h+/+ Tau+* and *Ch25h-/-Tau+* mice. H-I) Quantification of VGAT intensity and positive area in CA1 region of female *Ch25h+/+ Tau+* and *Ch25h-/-Tau+* mice. Unpaired student’s t-test. *p<0.05. n = 10 mice for *Ch25h+/+ Tau+*, n = 10 mice for *Ch25h-/-Tau+*. scale bar: 20µm. J) Representative western blot images of frontal cortical tissue from *Ch25h+/+ Tau+* and *Ch25h-/-Tau+* mice. K-L) Quantification of Synaptophysin levels and Psd95 levels of J. Unpaired student’s t-test: **p<0.01, ****p<0.0001. n = 12 mice for *Ch25h+/+ Tau+*, n = 12 mice for *Ch25h-/-Tau+*. M) Escape latency across the training days across *Ch25h+/+, Ch25h+/+ Tau+* and *Ch25h-/-Tau+* mice. Repeated-Measures Two-Way ANOVA. *p<0.05. N-O) Target annulus crossovers (N) and swimming speed (O) across genotypes. One-Way ANOVA followed by Bonferroni’s post hoc test. *p<0.05. n = 6 mice for *Ch25h+/+* mice, n = 7 mice for *Ch25h+/+ Tau+*, n = 6 mice for *Ch25h-/-Tau+*.

### Ch25h deficiency mitigates synapse loss and rescues spatial memory deficit in tauopathy mice

Next, we assessed the impact of *Ch25h* deletion on synaptic integrity. By staining the brain with antibody against VGAT, a well-characterized marker of presynaptic GABAergic and glycinergic neurons^50^, we found a significant increase of VGAT-positive synapses in the CA1 area of *Ch25h^-/-^ Tau^+^* mice compared to their littermates of *Ch25h^+/+^ Tau^+^* mice (**Fig. 1 G-I, supplementary dataset-1**). *Ch25h* deletion also significantly elevated the levels of pre-and post-synaptic proteins SYNAPTOPHYSIN and PSD95 in tauopathy mouse brains (**Fig. 1 J-L, supplementary dataset-1**).

To determine if *Ch25h* deletion affected tauopathy-induced spatial learning and memory deficits, we conducted the Morris water maze test. Spatial learning was assessed in the hidden platform trial over five consecutive training days by measuring the average latency to reach the platform each day. *Ch25h^+/+^ Tau^+^* mice exhibited significant impairments in spatial learning, whereas *Ch25h^-/-^ Tau^+^*mice showed improved performance comparable to *Ch25h^+/+^* wild-type controls (**Fig. 1M, supplementary dataset-1**). Twenty-four hours after the final training day, during the probe trials assessing spatial memory, *Tau^+^* mice made significant fewer target annulus crossovers than non-transgenic mice. Notably, *Ch25h* deficiency restored the frequency of target annulus crossings to levels similar to *Ch25h^+/+^*wild-type controls (**Fig. 1N, supplementary dataset-1**). Swimming speeds were comparable across all genotypes (**Fig. 1O, supplementary dataset-1**). Thus, *Ch25h* deletion protected against tau-induced synaptic loss and spatial learning and memory deficits.

### Ch25h deletion suppresses neuroinflammation and STING signaling in tauopathy mouse brains

To dissect the underlying mechanisms, we first examined how *Ch25h* deletion influences brain transcriptomes by performing bulk RNA-Seq in hippocampus. Differential gene expression analysis showed that *Ch25h* deletion reversed many tau-induced changes, including AD risk genes (*Apoe, Trem2, Grn, Clu, etc*) **(Fig. 2A**, **supplementary dataset-2**). Ingenuity Pathway analysis (IPA) revealed that these genes are enrichment in innate immune signaling (e. g., interferon, cytokine storm, TREM1, complement) and adaptive immune pathways (e.g. antigen presentation, T-cell activation/differentiation) (**Fig. 2B, supplementary dataset-2**). Heatmap for the expression of antigen presentation genes showed that *Ch25h* deletion significantly suppressed upregulation of both MHC class I and II by tau (**Supplementary Fig. 2A**), as confirmed by immunostaining (**Supplementary Fig. 2B-C, supplementary dataset-2**).

**Figure 2.**
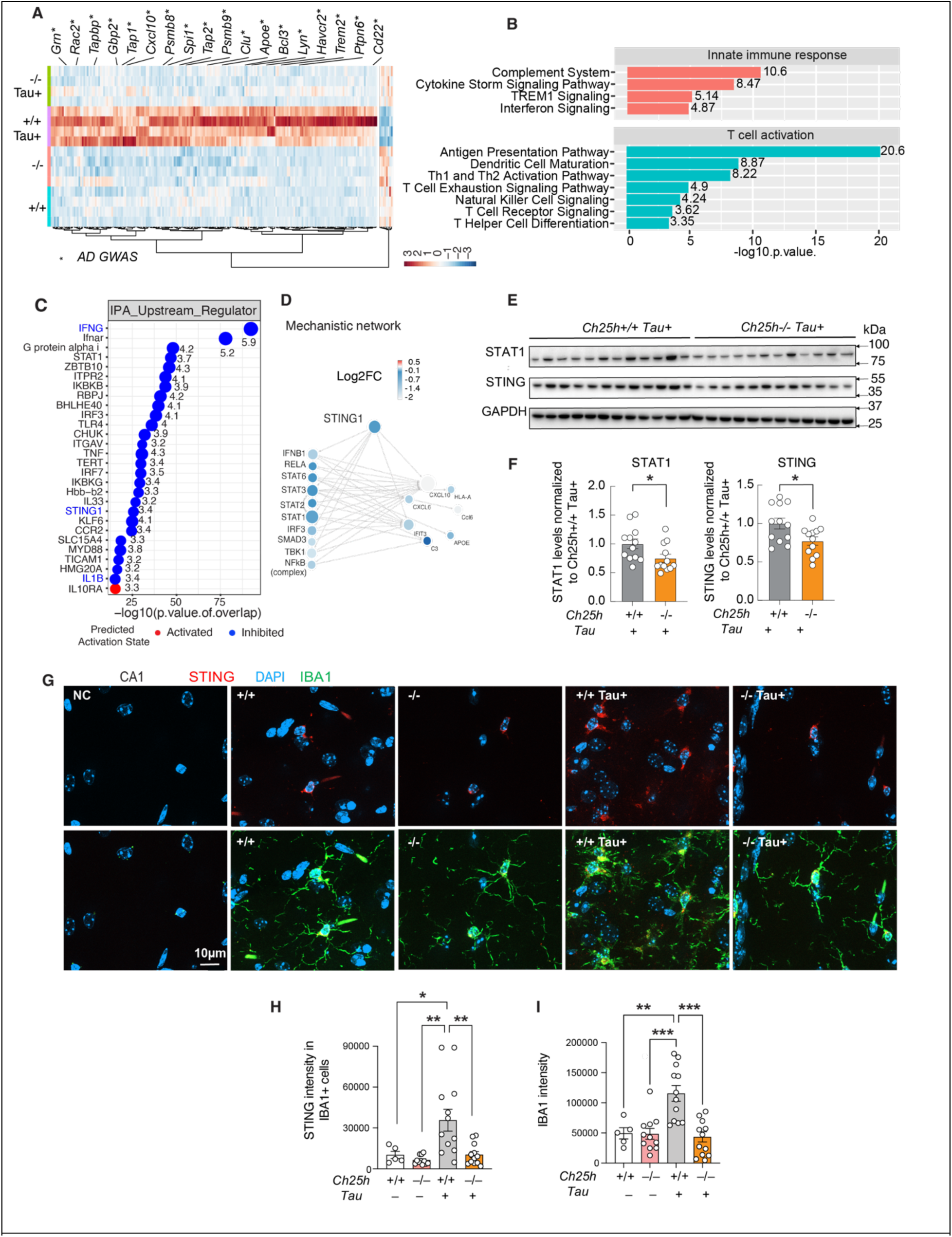
C*h*25h deletion suppresses microglial STING-IFN signaling in tauopathy mouse brain. A) Heatmap comparison of DEGs genes increased in *Ch25h+/+ Tau+* mice and reversed by *Ch25h* deletion. Asterisk represents AD GWAS genes. B) Enriched pathways predicted by GSEA for 199 DEGs upregulated in *Ch25h+/+ Tau+* mice and reversed in *Ch25h-/-Tau+* mice. C) Upstream regulators predicted by IPA for 199 genes increased in *Ch25h+/+ Tau+* mice and reversed by *Ch25h* deletion. D) Mechanistic network of STING predicted by IPA upstream regulator analysis in (C). E) Representative immunoblot images of STAT1 and STING in *Ch25h+/+ Tau+* and *Ch25h-/-Tau+* mice. F) Quantification of STAT1 and STING in *Ch25h+/+ Tau+* and *Ch25h-/-Tau+* mice. n = 12 mice for *Ch25h+/+ Tau+*, n = 12 mice for *Ch25h-/-Tau+.* Unpaired student’s t-test. *p<0.05. G) Representative images showing STING and IBA1 staining. NC: negative control without primary antibodies. H–I) Quantification (H–I) of the STING-positive area within IBA1-positive cells and IBA1 intensity in the CA1 region of the hippocampus. Two-Way ANOVA, *p<0.05, **p<0.01, ***p<0.001. n = 5 mice for *Ch25h+/+ Tau-* mice, n = 11 mice for *Ch25h-/-Tau-,* n = 12 mice for *Ch25h+/+ Tau+*, n = 11 mice for *Ch25h-/-Tau+*.

To identify the upstream regulators responsible for these changes, we performed IPA upstream regulators analysis and identified top regulators including IFNG, IFNAR, STING1, and STAT1 that are inhibited by *Ch25h* deletion (**Fig. 2C, supplementary dataset-2**). Network analysis highlights a suppression of STING-mediated proinflammatory signaling by *Ch25h* deletion (**Fig. 2D**). Western blot revealed that *Ch25h* deletion reduced STING and STAT1 protein levels in brain (**Fig. 2E-F, supplementary dataset-2**). Co-immunostaining of STING and IBA1 demonstrated that an increase of microglial STING signals in CA1 region in *Ch25h ^+/+^ Tau^+^* mice was suppressed by *Ch25h* deletion (**Fig. 2G-H, supplementary dataset-2**), accompanied with a reduction of IBA1 intensity (**Fig. 2I, supplementary dataset-2**).

### Ch25h amplifies STING IFN signaling through a positive feedback loop in vivo

To directly examine how CH25H regulates microglial STING activation, we treated primary microglia with the STING agonist DMXAA, which drives STING translocation from ER to Golgi, a crucial step in STING signaling and activation^51^. Co-staining for STING and the Golgi marker GM130 revealed that deletion of Ch25h significantly reduced the amount of STING localized in Golgi (**Fig. 3A-C, supplementary dataset-3**). Western blot further confirmed that DMXAA-induced phospho-STING (Ser365) upregulation was suppressed by *Ch25h* deficiency (**Fig. 3D-E, supplementary dataset-3**). These results suggest that CH25H is required for STING trafficking and activation in microglia.

**Figure 3.**
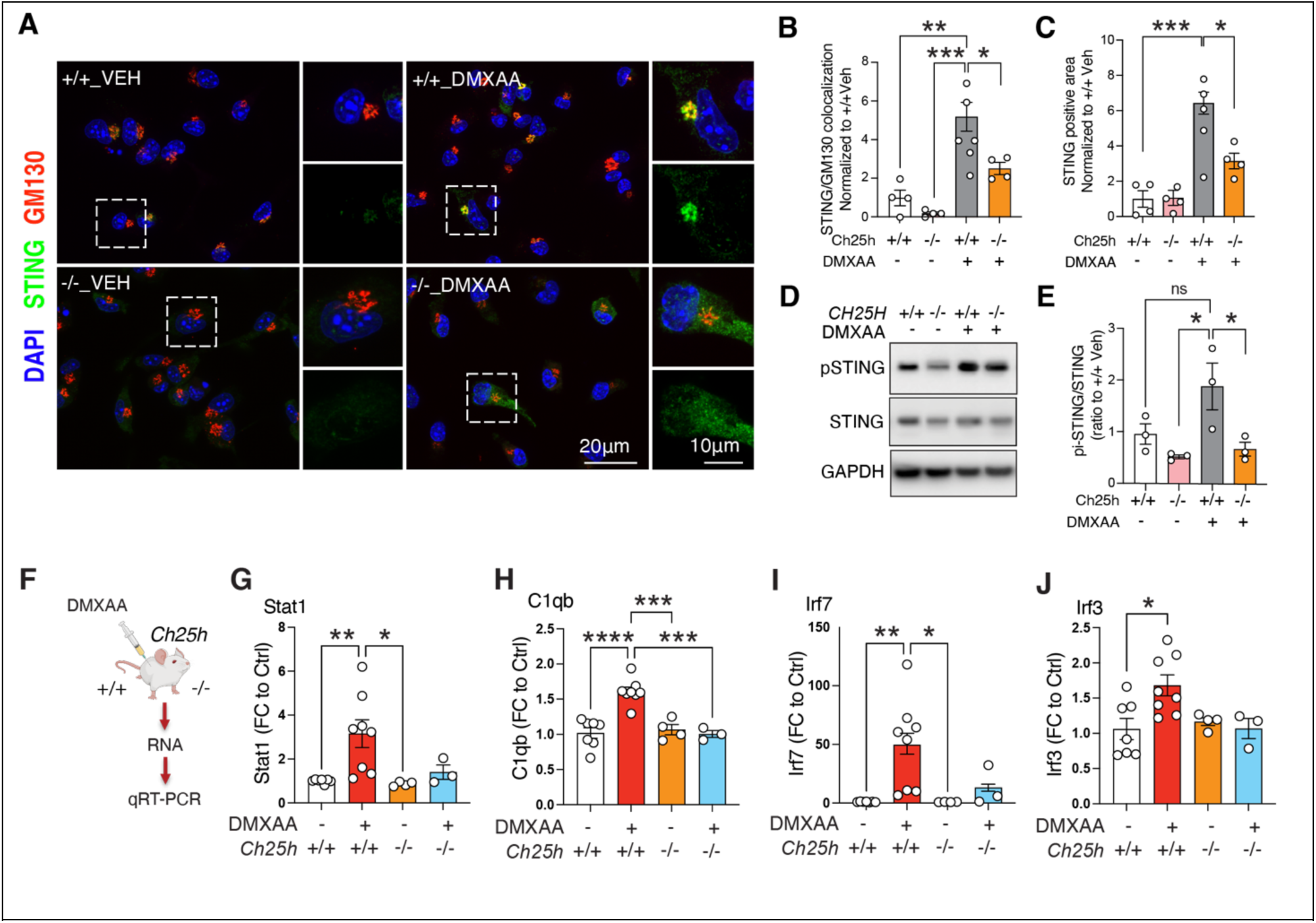
C*h*25h deletion suppresses microglial STING translocation to Golgi and blocks IFN response in vivo. A) Representative images of immunostaining using STING and GM130 antibodies in primary microglia treated with DMSO vehicle or DMXAA for 4hrs. B) Quantification of STING and GM130 colocalized area. C) Quantification of total STING positive area. Data points from three independent experiments (126 cells from *Ch25h^+/+^,* 108 cells from *Ch25h^-/-^,* 136 cells from *Ch25h^+/+^*DMXAA, 130 cells from *Ch25h^-/-^* DMXAA). D) Representative immunoblot images of pSTING (Ser365) and STING in Ch25h+/+ or Ch25h-/-primary mouse microglia treated with or without DMXAA. GAPDH as internal control. E) Quantification of pi-STING(Ser365) and total STING normalized by GAPDH based on 3 independent experiments. F) Diagram of in vivo DMXAA treatment. *Ch25h^+/+^* and *Ch25h^-/-^* mice were i.p. injected with vehicle or 25 mg/kg DMXAA for 6hrs followed with brain dissection and RNA extraction. The levels of IFN genes were analyzed by quantitative RT-PCR. G-J) Quantification of Stat1, C1qb, Irf7 and Irf3 mRNA across four treatment conditions. Each data point represents one mouse sample. Statistical analyses in B, C, E, G-J were performed by Two-Way ANOVA followed by Tukey post-hoc test. *p<0.05, **p<0.01, ***p<0.001, ****p<0.0001.

We then asked whether this requirement extends to IFN signaling in vivo. Following DMXAA administration in WT and *Ch25h* deficient mice (**Fig. 3F**), qPCR analysis of hippocampal tissue showed that the significant inductions of Stat1, C1qb, Ifn3 and Ifn7 by DMXAA were no longer significant in the absence of *Ch25h* (**Fig. 3G-J, supplementary dataset-3**).

To understand how IFN signaling in turn affects Ch25h, we reanalyzed our prior snRNA seq dataset^52^. DMXAA increased *Ch25h* expression in microglia in WT mice, but this induction was abolished in IFNAR deficient mice (**Supplementary Fig. 3A, B**). A parallel experiment in WT and STING KO mice showed the same requirement: *Sting1* deficiency prevented *Ch25h* upregulation (**Supplementary Fig. 3A, C**), further supporting that IFN signaling promotes *Ch25h* expression in microglia. Together, our results reveal a positive feedback loop in which IFN activation induces *Ch25h*, and *Ch25h* in turn enhances STING-IFN pathway activation.

### Ch25h deletion restores homeostatic microglial gene expression in tauopathy mice

To investigate the effects of *Ch25h* deletion at single cell level in vivo, we performed single nuclei RNA sequencing (snRNA-Seq) of hippocampal tissues across the four genotypes. After quality control and removal of multiplets using DoubletFinder ^53^ **(Supplementary Fig. 4A-D**), 71,716 high-quality nuclei were clustered into seven major brain cell types (**Supplementary Fig. 5A-C**). Differential gene expression analyses demonstrate that excitatory neurons (EN) and astrocytes (AST) have the highest numbers of DEGs (**Supplementary Fig. 5D**).

*Ch25h* expression predominantly in the microglia, with minimal expression in vascular cells (**Supplementary Fig. 5E**). Pseudobulk DEG analysis of microglial cluster showed that *Ch25h* deletion restored homeostatic microglial genes (*P2ry12, Cx3cr1, Hexb, Cst3, csf1r, Tmem119*) (**Fig. 4A**) and suppressed tau-induced upregulation of DAM genes including *Apoe, Clec7a,* and *Cd74* (**Fig. 4B**). Subcluster analysis of microglia identified three subclusters (**Supplementary Fig. 5F**). Tau significantly expanded the size of subcluster-2. Tau also reduced subgroup-1, which was restored by *Ch25h* deletion. Correlation analysis showed that subgroup-2 is more DAM-like while subcluster 1 is more homeostatic (**Supplementary Fig. 5G-H**). Immunostaining validated that P2RY12 levels are significantly increased in CA1 microglia of *Ch25h^-/-^ Tau^+^* mice compared to *Ch25h^+/+^ Tau^+^* littermates (**Fig. 4C-D, supplementary dataset-4**). These results suggest a shift of microglia from disease-associated to homeostatic state in tauopathy mouse brains in the absence of *Ch25h*.

**Figure 4.**
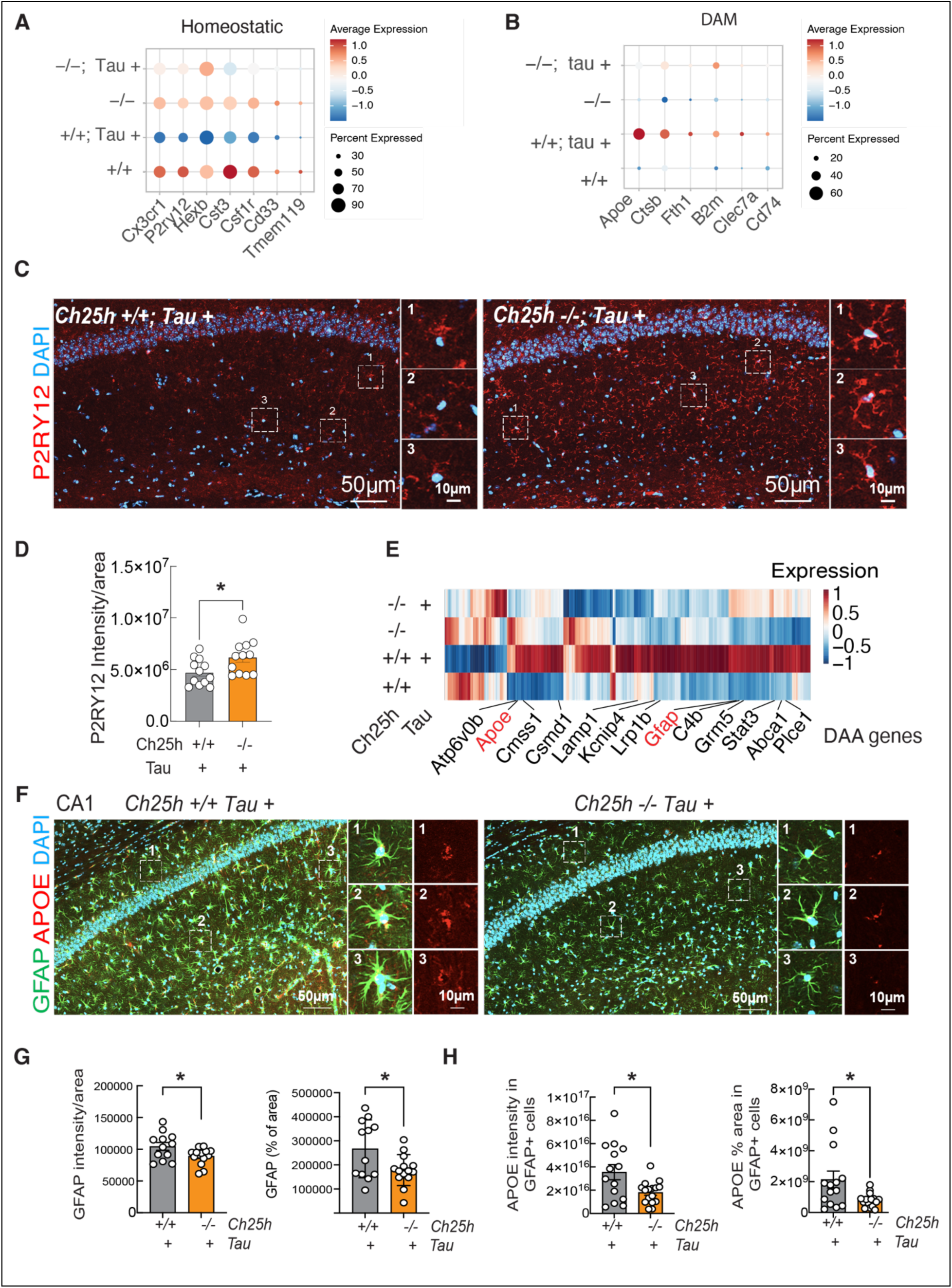
*Ch25h* deletion suppresses disease associated astrocytic genes and reduces astrocytic APOE levels. A-B) Dot plot of expressions of selective homeostatic (A) or disease-associated microglial (DAM) genes (B) in snRNAseq data from 9-month-old *Ch25h+/+ Tau-, Ch25h-/-Tau-, Ch25h+/+ Tau+,* and *Ch25h-/-Tau+* mice. n=4 per genotype. C) Representative images of P2RY12 staining across four genotypes. D) Quantification of microglial P2RY12 intensity in CA1 region of *Ch25h+/+ Tau+,* and *Ch25h-/-Tau+* mice. Scale bar: 50μm. Insert: representative microglia. Scale bar: 10μm. Unpaired student’s t-test: *p<0.05. n=12 per genotype. E) Heatmap comparison of genes differentially expressed in astrocyte across four genotypes. Log2FC ≥ 0.1 or ≤-0.1, FDR£0.05. Previously identified CAA (Disease-Associated Astrocytes) genes were shown (Habib, et al, 2020). F) Representative images of GFAP (green) and APOE (red) co-staining in CA1 region of *Ch25h+/+ Tau+,* and *Ch25h-/-Tau+* mice. G) Quantification of total GFAP intensity or positive area in CA1 region of *Ch25h+/+ Tau+,* and *Ch25h-/-Tau+* mice H) Quantification of APOE intensity or positive area in GFAP positive cells in CA1 region of *Ch25h+/+ Tau+,* and *Ch25h-/-Tau+* mice. Scale bar: 50μm. Insert: three representative microglia. Scale bar: 10μm, Unpaired student’s t-test: *p<0.05. n=12 mice per genotype.

### Ch25h deficiency represses expression of disease-associated astrocytic genes in tauopathy mice

We next examined the effect of *Ch25h* deficiency on astrocytes, which were found to be regulated by *Ch25h* and 25-HC recently^47,54^. Pseudobulk analysis revealed that *Ch25h* deletion reversed 262 DEGs altered by tau (**Fig. 4E**), with 218 up by tau and 44 down by tau (**Supplementary Fig. 6A-B**, supplementary table 5). Nineteen of them are overlapped with disease-associated astrocyte (DAA) genes previously identified in AD and aging^55^, including *Apoe, Clu, Abca1, Gfap, Lrp1b, C4b,* and *Stat3* (**Fig. 4E**). GSEA analysis of these 262 DEGs revealed that they are enriched in multiple cellular pathways (**Supplementary Fig. 6C-D**). Notably, the top pathway suppressed by Tau but reserved by *Ch25h* deletion is cholesterol homeostasis (**Supplementary Fig. 6D).** Dot plot further showed that Tau markedly suppressed multiple key regulators of de novo cholesterol biosynthesis, which were normalized by *Ch25h* deletion (**supplementary Fig. 6E**). IPA analyses of DEGs reversed by *Ch25h* deletion also highlighted LXR/RXR signaling with genes involved in cholesterol transport such as Abca1, Apoe, and Clu (**Supplementary Fig. 6F**). Immunostaining demonstrated a reduction of GFAP intensity and morphological change from highly branched hypertrophic astrocytes in *Ch25h^+/+^Tau^+^* mice toward simple star-shaped in *Ch25h^-/-^Tau^+^* mice, along with significantly lower GFAP levels and less ApoE in GFAP+ astrocytes (**Fig.4F-H, supplementary dataset-4**). Collectively, these results suggest that CH25H mediates tau-induced disruption of astrocytic cholesterol metabolism.

### Ch25h mediates cholesterol esters accumulate in PS19 tauopathy mouse brain

To further dissect how CH25H influences lipid metabolism in tauopathy, we performed lipidomic profiling using liquid chromatography-tandem mass spectrometry (LC-MS/MS). Lipidomic analyses revealed a marked increase in total cholesterol esters (CEs) in the cortices of 9-month-old *P301S Tau+* mice compared to non-transgenic littermates (**Fig. 5A, supplementary dataset-5**). The proportion of CEs within the total lipid composition rose dramatically from 0.99% in the brains of control mice (Tau-) to 16.05% in Tau+ brains (**Fig. 5B**), while the total free cholesterol (FC) levels remained unchanged (**Fig. 5A**). Analyses of individual CE subspecies showed broad elevation across most of CE species in Tau+ brains (**Fig. 5C)**. *Ch25h* deletion significantly suppressed the tau-induced elevation of CEs (**Fig. 5C**), demonstrating a critical role for CH25H in promoting CE accumulation in tauopathy.

**Figure 5.**
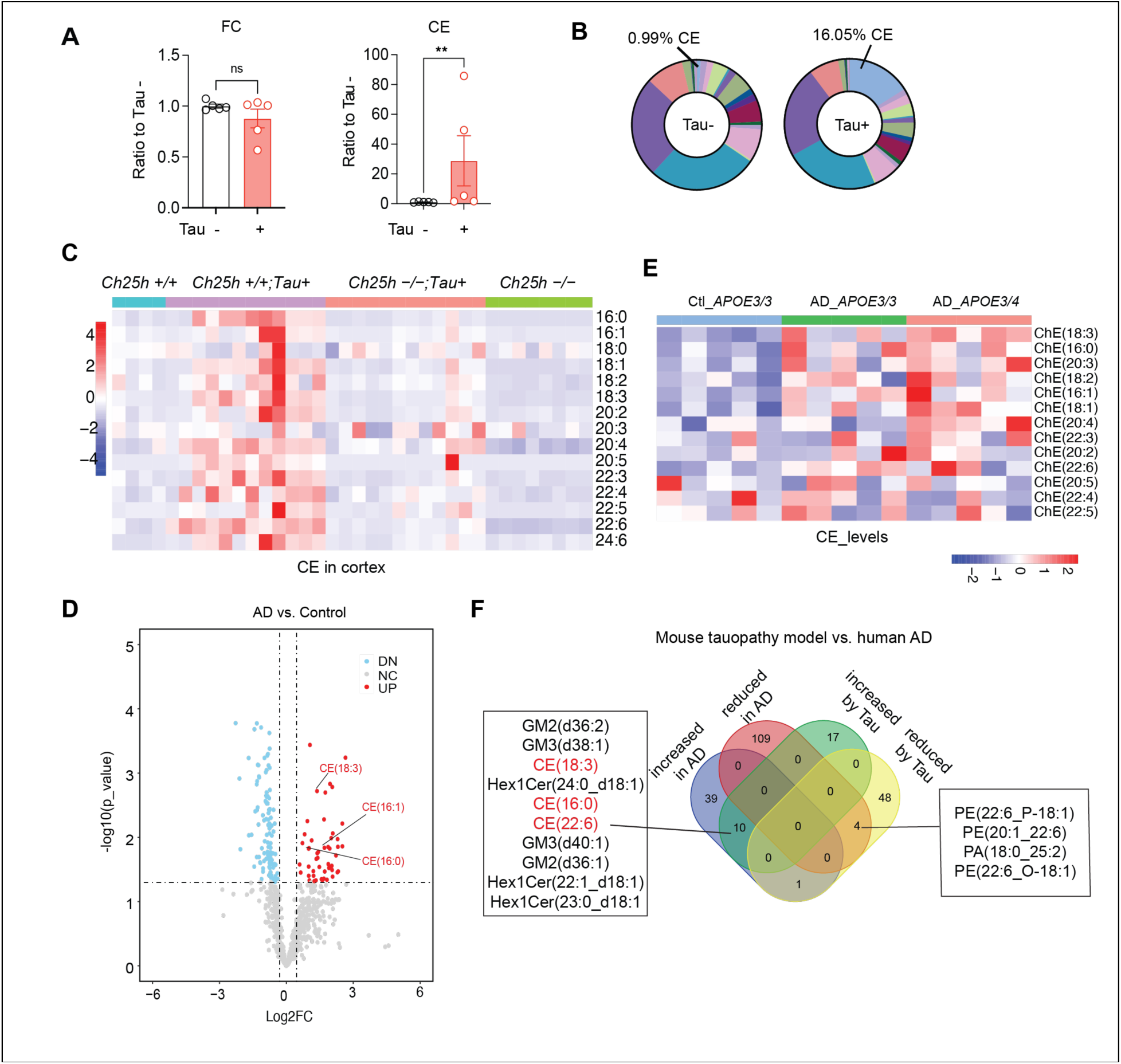
Ch25h regulates cholesterol ester accumulation in PS19 tauopathy mouse brains. A) Quantification of free cholesterol (FC, left) and cholesterol esters (CE, right) in cortical tissues from *Tau+* mice and non-transgenic littermates (*Tau-*). N=5/genotype. non-parametric Mann-Whitney U test, **p<0.01. B) Relative composition of CE species vs total lipids in cortical tissues of *Tau−* and *Tau+* mice. C) Heatmap showing levels of CE species in cortical tissues of female mice across genotypes (n=4 mice for *Ch25h+/+,* n=8 mice for *Ch25h−/−,* n=12 mice for *Ch25h+/+;Tau*+, n=12 mice for *Ch25h−/−;Tau+*). D) Volcano plot of lipids altered in human AD brains compared to matched non-dementia controls. N=5 for control, N=10 for AD, all females. E) Heatmap showing the levels CE species in AD and control brains. F) Overlapped lipid species altered in human AD and tauopathy mouse brains.

Consistent with these, lipidomic analyses in human brain revealed that several CE species were significantly elevated in the frontal cortical tissue of AD brains compared to non-dementia controls, with a trend toward higher levels in APOE4 carriers (**Fig. 5D-E**). Comparative analysis between human and mouse lipidomic datasets identified shared lipid perturbations, including CE 18:3; CE 16:0, and CE 22:6, which were upregulated in both human AD and tauopathy mouse brains but restored by *Ch25h* deletion (**Fig. 5C, F**).

### 25-HC reprograms neuronal lipids and induces cellular stress in human neurons

We next directly examined how 25-HC affects lipid profiles in human tauopathy neurons, which are engineered to express 4R-P301S tau and develop robust MC1 positive tau inclusion when seeded with K18 fibrils^56^. Human 4R-P301S iPSC were differentiated into neurons and exposed to tau seeds together with 25-HC for 14 days (**supplementary Fig. 7A**). Using LC-MS/MS, we quantified sterol lipids, glycerolipids, glycerophospholipids, and sphingolipids, four major lipid classes involved in energy homeostasis, membrane integrity, and lipid signaling^57–61^. Tau seeds alone caused to a mild reduction of some lipid classes (**supplementary Fig. 7B**). Strikingly, 25- HC profoundly induced alterations in many lipid species **(supplementary Fig. 7C-E).** Notably, CEs were significantly increased following 25-HC exposure (**Fig. 6 A, B, supplementary dataset-6**). Furthermore, 25-HC exacerbated the tau-seed induced loss of lysosomal lipid bis(monoacylglycerol)phosphates (BMPs) ^59^, while selectively increasing BMP 40:7, suggesting a shift of lysosomal BMP composition (**Fig. 6C, D, supplementary dataset-6**). Lipidomics also revealed a dramatic decrease of cardiolipin (CL), a phospholipid confined to the inner mitochondrial membrane and critical for mitochondrial function^62,63^ (**Fig. 6E, supplementary dataset-6**), indicative of mitochondrial dysfunction and stress. Profiling of phosphotidylglycerol (PG), a biosynthetic precursor for both lysosomal BMP^59,64^ and mitochondrial CL^65^(**Fig. 6F**), showed a global reduction of total PG (**Fig. 6G, supplementary dataset-6**), but a selective enrichment of long-chain polyunsaturated PG species (**supplementary Fig. 7F**), consistent with compensatory membrane lipid remodeling associate with mitochondrial or lysosomal membrane. In contrast, phosphatidic acid (PA), a key precursor for PG, CL, and BMP^59,66^ (**Fig. 6F**), was significantly elevated upon 25-HC or 25-HC/tau seeds treatment (**Fig. 6H, I, supplementary dataset-6**), suggesting a metabolic bottleneck in PA conversion into downstream glycerophospholipids and a disruption of the PG-CL or PG-BMP axis.

**Figure 6.**
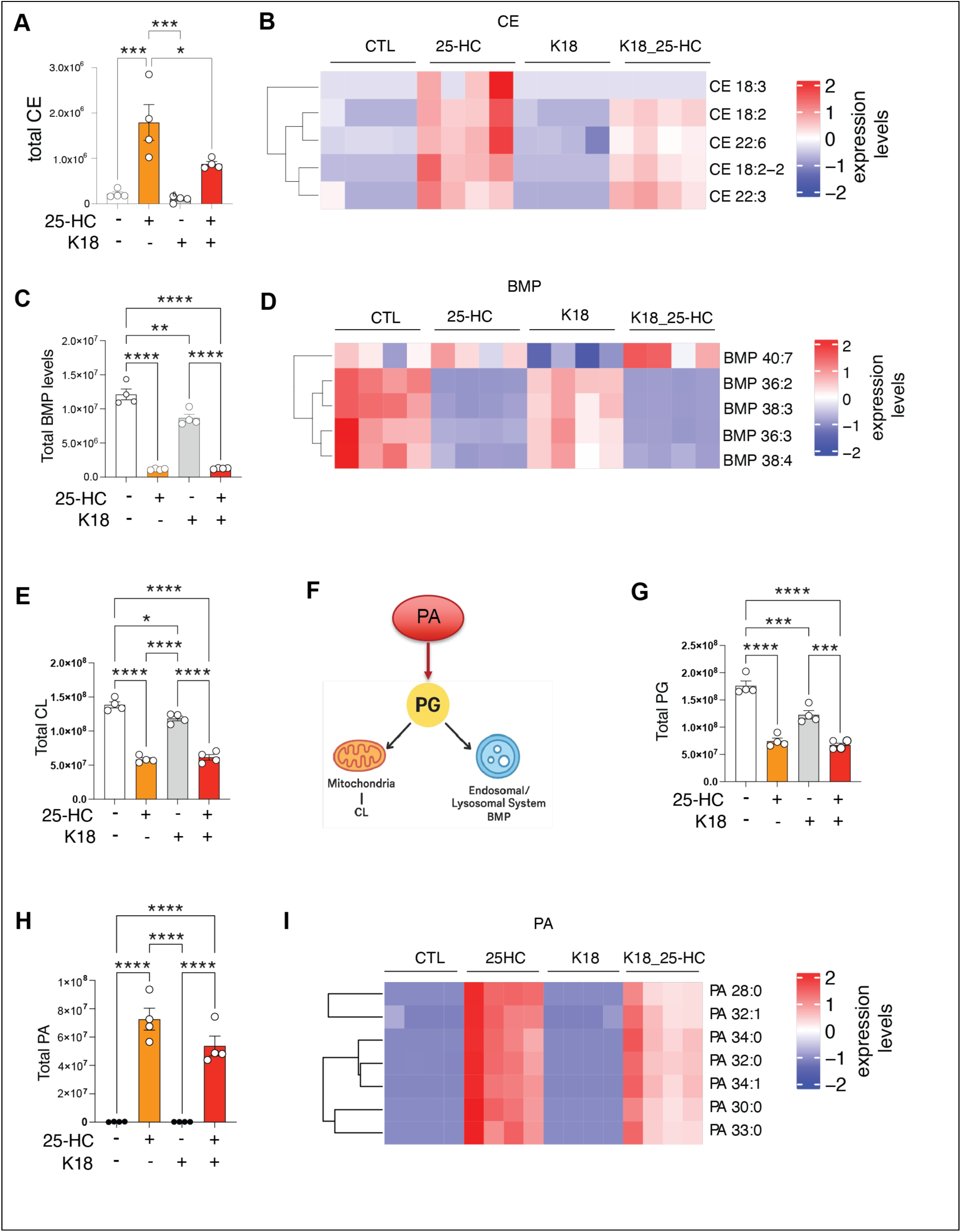
25-HC reprograms lipid metabolism in human neurons. Lipid species altered by tau seeds and 25-HC in human 4R-P301S neurons. Neurons were exposed to 1.5 mg/mL K18 tau fibrils with or without 25-HC at day 1 and maintained in culture for 14 days. Lipidomic analysis in human 4R-P301S neurons across four treatment conditions: A) Total CE levels. B) Heatmap of CE species. C) Total BMP levels. D) Heatmap of BMP species. E) Total Cardiolipin levels. F) Diagram of metabolism relationships among PA, PG, CL and BMP. G) Quantification of total PG. H) Quantification of total PA. I) Heatmap of PA species. Statistical analyses in A, C, E, G, H were performed by Two-Way ANOVA followed by Tukey post-hoc test. *p<0.05, **p<0.01, ***p<0.001, ****p<0.0001.

We also detected a marked increase in lysophosphatidylcholine (LPC) lipids, particularly those short chain, bioactive species known for proinflammatory activity^67–70^ (**supplementary Fig. 7G-H**), indicating enhanced membrane turnover, oxidative stress, or inflammation. Additional lipids significantly decreased include Sphingomyelin (SM), Hexosylceramide (HexCer), Lysophosphatidylinositol (LPI) and Lysophosphatidylglycerol (LPG) (**supplementary Fig. 7I**).

### 25-HC promotes tau propagation in human tauopathy neurons

The profoundly lysosomal and mitochondrial stress resulted from reprograming of lipid metabolism in human neurons in combination of marked reduction of tau pathology by *Ch25h* deletion in vivo prompted us to test whether microglial-derived 25-HC directly influences tau propagation in K18 seeded 4R-P301S human neurons^56^ (**Fig. 7A**). Tau solubility was assessed by sequential extraction with Triton X and Sodium Dodecyl Sulfate (SDS). Quantitative western blot analysis revealed that 25-HC treatment markedly enhanced Triton-insoluble tau levels induced by tau seeds, shifting tau from soluble to insoluble fraction (**Fig. 7B-D, supplementary dataset-7**). This effect was dose-dependent and specific to 25-HC, as cholesterol treatment had no measurable impact on tau inclusions (**Fig. 7E-F, supplementary dataset-7**). In parallel, co-treatment of 25- HC with tau seeds for two weeks significantly increased MC1+ tau inclusions (**Fig. 7G-H, supplementary dataset-7**).

**Figure 7:**
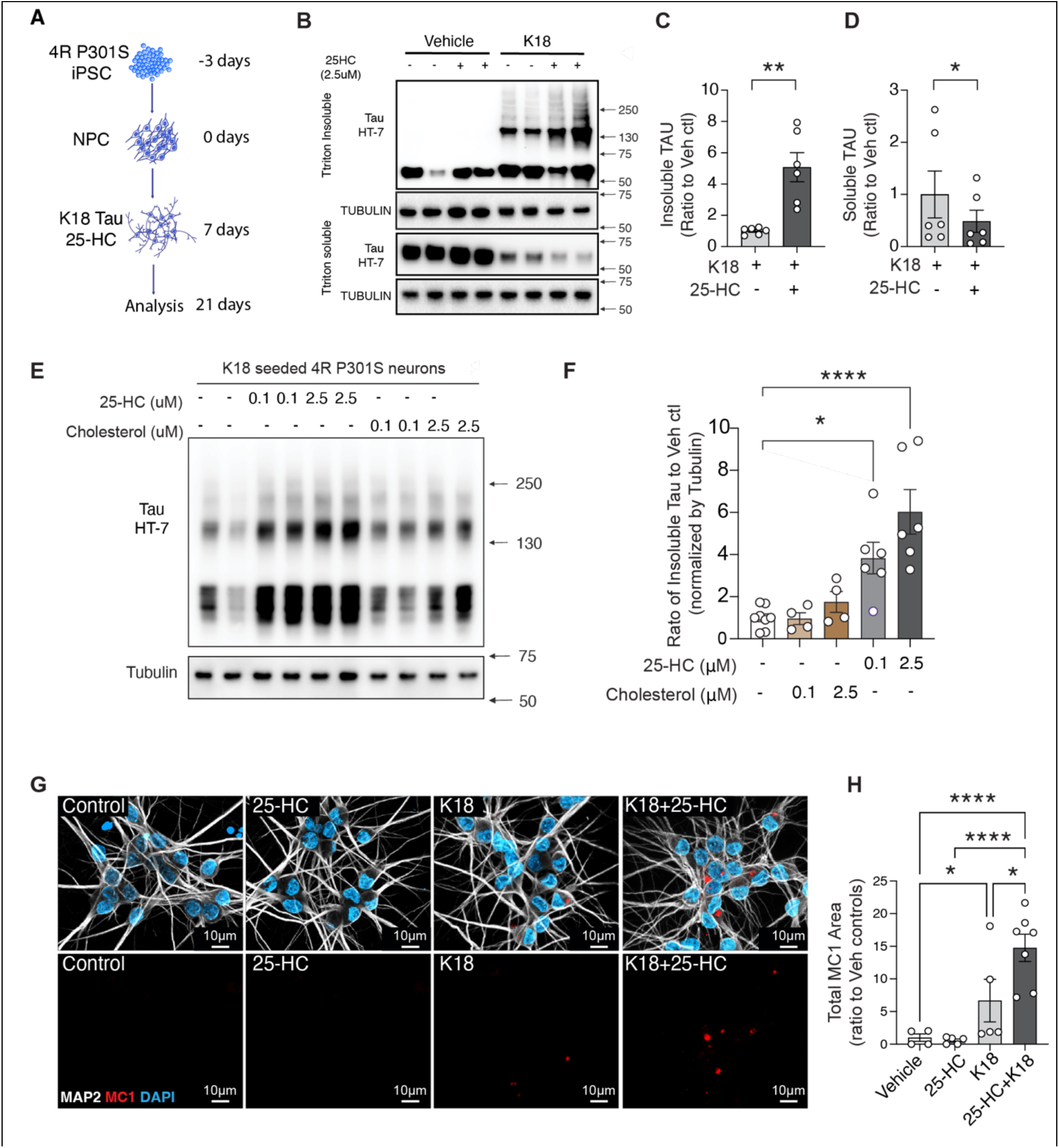
25-HC exaggerates tau aggregation in human IPSC derived neurons. A) Schematic for human iPSC-derived neuron differentiation and treatment. B) Representative immunoblot images of triton soluble and insoluble fractionations from D28 (7 +21) 4R-P301S neurons ±1.5 mg/mL K18 seeding, and with ± 2.5uM 25-HC treatment. C-D) Quantification of HT-7 positive tau in insoluble fractions (C) or soluble fraction (D) after normalization to TUBULIN. *p < 0.05, **p < 0.01, unpaired student test. Data points were obtained from three independent experiments. E) Representative immunoblot images of Triton-insoluble lysates from D28 (7 +21) 4R-P301S neurons treated with K18 and 2.5µM 25-HC or 2.5µM Cholesterol. F) Quantification of HT-7 positive tau in Triton-insoluble fractions normalized to TUBULIN. *p < 0.05, One-Way ANOVA followed by Tukey post-hoc test from three independent experiments. G) Representative images of MC1 positive tau and MAP2 staining across four conditions. Scale bar, 50μm. H) Quantification of (G). Data points were from three independent experiments. *p<0.05, ****p<0.0001. n = 4 wells for Vehicle, n = 5 wells for 25-HC, n = 5 wells for K18, n = 7 wells for K18+25-HC. n = 5 images per well.

### 25-HC promotes tauopathy-induced apoptosis in human neurons

Deletion of *Ch25h* markedly ameliorated tau-driven neurodegeneration and reduced tau pathology. To evaluate whether 25-HC directly contributes to neuronal injury, we examined human 4R P301S neurons treated with K18 tau fibrils and 25-HC using TUNEL staining, which detects programmed cell death (apoptosis) by labeling fragmented DNA. Although 25-HC alone caused modest toxicity in both 4R P301S and 4R WT neurons, K18 induced apoptosis only in 4R P301S neurons, where MC1 positive tau inclusions formed, and not in 4R WT neurons lacking inclusions, and this toxic effect of K18 was further intensified by 25-HC (**Fig. 8A-D, supplementary dataset-8**). Moreover, both agents sharply reduced the synaptic proteins PSD95 and SYNAPTOPHYSIN (**Fig. 8E-G, supplementary dataset-8**), indicating loss of synaptic integrity and demonstrating synergistic toxicity from tau seeds and 25-HC. Together, these findings identify 25-HC, a microglial derived lipid mediator, as a direct amplifier of tau pathology and tau-induced neurodegeneration.

**Figure 8.**
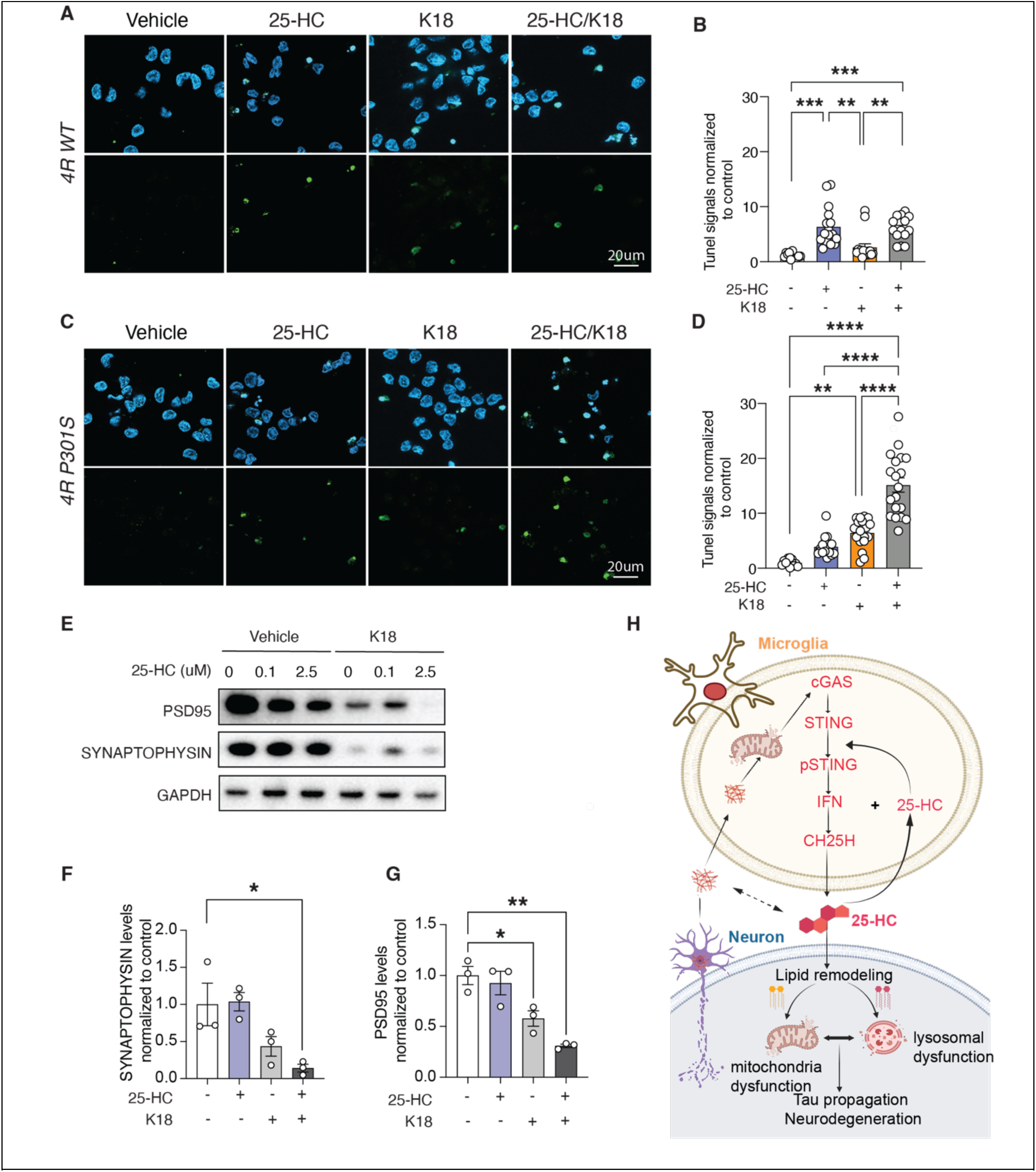
25-HC reduces synaptic markers and induces apoptosis in human neurons. A–B) Representative images (A) and quantification (B) of TUNEL Staining for 4R human neurons under different treatments. Three independent experiments. **p < 0.01, ***p < 0.001. Two-Way ANOVA followed by Tukey post-hoc test. C–D) Representative images (C) and quantification (D) of TUNEL Staining for 4R P301S human neurons under different treatments. Three independent experiments **p < 0.01. ****p < 0.0001. Two-Way ANOVA followed by Tukey post-hoc test. E) Representative immunoblot images of PDS95 and Synaptophysin in human 4R-P301S neurons seeded with K18 tau in the presence of 25-HC vs controls. F–G) Quantification of PSD95 and Synaptophysin levels. Three independent experiments. *p < 0.05, **p < 0.01. TWO-WAY ANOVA followed by Tukey post-hoc test. H) Diagram illustrating the proposed mechanisms through which CH25H and 25-HC act via the STING–IFN–CH25H lipid axis to bridge innate immune activation to tau pathology and toxicity in AD and related tauopathies.

## Discussion

Our study identifies CH25H as a critical effector and modulator of microglial STING-IFN signaling in the neuroinflammatory and neurodegenerative cascades driven by tau pathology. We demonstrate that genetic deletion of *Ch25h* in female *P301S* tauopathy mice markedly reduces tau accumulation, preserves synaptic integrity, and improves cognitive performance. Mechanistically, IFN activation upregulates *Ch25h* in microglia, while *Ch25h* loss suppresses microglial STING– interferon signaling, restores homeostatic glial transcriptional programs, and prevents lipid dysregulation in brain. In contrast, the oxysterol product 25-HC directly promotes neuronal tau aggregation, synaptic loss, and apoptosis through profound remodeling of neuronal lipid metabolism. Together, these findings reveal a feedforward STING–CH25H–25-HC axis linking microglial innate immune activation to tau propagation and tau driven neurotoxicity.

Microglia in neurodegenerative brains exhibit a disease-associated phenotype accompanied by broad lipid metabolic reprogramming^11,12,16,17,19,71,72^. IFN signaling is one of the key factors driven tau toxicity and tauopathy^52,71,73–76^. Our transcriptomic data show that Ch25h deletion reverses many of these changes, including suppression of *Apoe, Clec7a*, and *Cd74* while restoring homeostatic markers such as *P2ry12* and *Cx3cr1*. Network analyses identify *STING* and *STAT1* as upstream regulators inhibited by *Ch25h* loss, and our in vitro imaging and biochemical results demonstrate that *Ch25h*-deficient microglia fail to traffic STING to the Golgi and exhibit reduced STING phosphorylation upon activation. These findings suggest that CH25H and its product 25- HC sustain microglial interferon signaling by facilitating STING activation.

Recent studies demonstrate an intimate crosstalk between cellular cholesterol and IFN signaling. York et al identified a cholesterol metabolism-type I interferon (IFN) inflammatory circuit, where interferon reprogram cholesterol homeostasis and perturbation of cholesterol biosynthesis promotes type I IFN responses in macrophages and T cells^77^. Refined studies further revealed that activation of STING, a critical player in IFN signaling, are tightly controlled at ER-Golgi interface^78–83^. Notably, cholesterol is one of the lipid regulators for STING ER exit ^77,81,84,85^. STING contains cholesterol-binding motifs^84^, which are critical for its ER retention and activation in Golgi^84,85^. Decline of cellular cholesterol promotes STING exit from ER, a critical step for its activation, likely via a mechanism reshaping ER membrane curvature^84^. 25-HC is a critical regulator for cholesterol metabolism by inhibiting SREBP2 activity in elevating cellular cholesterol biosynthesis^27,86^. Thus, reprograming cellular cholesterol by 25-HC may provide a lipid environment that supports persistent STING signaling. Our study revealed a bidirectional regulation between STING signaling and lipid metabolism. STING-IFN activation upregulates CH25H, which metabolizes cholesterol to 25-HC, and this 25-HC in turn promotes STING signaling, creating a positive feedback loop that contributes to the sustained STING-IFN activation in tauopathy.

Emerging studies suggest that glia-derived lipids can act as diffusible mediators of neurotoxicity beyond classical cytokine pathways ^11,87,88^. Using a human iPSC-derived tauopathy neuron model, we find that 25-HC directly enhances seeded tau aggregation and triggers neuronal apoptosis, positioning it as a key downstream effector of microglial STING-IFN signaling. Lipidomic profiling further reveals that 25-HC drives cholesterol ester accumulation, reduces lysosomal BMP, and mitochondrial cardiolipin, changes that collectively induce lysosomal and mitochondrial stress and compromise their functions, thereby promoting tau aggregation and cell death. These findings align with lipid abnormalities widely reported in AD^89–97^, including disrupted BMP levels in human and mouse AD brains^59,90,98,99^ and AD-associated PLD3 mutation that impair BMP production^100–103^. The 25-HC-induced loss of BMP, selective rise in BMP 40:7, and increase in neuronal CEs indicate exacerbated lysosomal stress and impaired degradative capability, favoring pathological tau accumulation. Likewise, the marked reduction of cardiolipin, previously linked to early mitochondrial defects in AD^104,105^, together with decreases in PG and other phospholipids and elevation of PA, suggests a metabolic bottleneck in the PG-BMP and PG-CL pathways. The accumulation of short chain LPCs, proinflammatory lipids enriched in AD brains^68,106,107^, further reflects heightened oxidative and inflammatory stress. Together, these data support a model in which IFN-induced 25-HC synergizes with tau seeds to drive a feed-forward loop of lysosomal and mitochondrial dysfunction, neuroinflammation and tau propagation in AD. Our single-nucleus analyses reveal that Ch25h deletion also reprograms astrocytes toward a more homeostatic transcriptional state, reversing the expression of multiple disease-associated astrocytic (DAA) genes, including *Apoe*, *Clu*, *Abca1*, and *Gfap*. These changes coincide with restoration of cholesterol biosynthetic and LXR/RXR pathways, suggesting that upregulation of microglial CH25H by tauopathy suppresses astrocytic lipid metabolism through paracrine oxysterol signaling. Prior studies have shown that 25-HC inhibits SREBP2 activation and limits de novo cholesterol synthesis, potentially accounting for the astrocytic lipid deficits observed in tauopathy^54^. By restoring astrocytic cholesterol homeostasis, Ch25h deletion may help preserve neuronal membrane maintenance and synaptic stability. These data support a model in which CH25H-driven lipid imbalance in microglia propagates metabolic dysfunction across glial networks.

In summary, this study delineates a mechanistic framework in which microglial CH25H and 25-HC couple STING-dependent innate immunity to lipid metabolic stress and tau-mediated neurodegeneration. By integrating immune and metabolic signaling across microglia, astrocytes, and neurons, CH25H functions as a central amplifier of neuroinflammatory injury. Several important questions remain to be addressed in future, including the mechanisms by which CH25H/25-HC promote STING trafficking, and how 25-HC–induced lipid remodeling influences neuronal tau aggregation and survival. From a translational perspective, pharmacologic inhibition of CH25H or blockade of 25-HC synthesis could interrupt the inflammatory–lipid feedback loop that sustains neurodegeneration. Targeting the STING–CH25H–25-HC axis may therefore represent a promising therapeutic strategy for Alzheimer’s disease and related tauopathies.

## Materials and Methods

### Animals

Mice were housed in groups up to five per cage with ad libitum access to food and water. All animals were maintained in a specific pathogen-free barrier facility under controlled conditions at a temperature of 21–23 °C, humidity ranging from 30% to 70%, and a 12-hour light/12-hour dark cycle. Human *PS19 Tau* transgenic mice expressing the T34 isoform of microtubule-associated protein tau with one N-terminal insert and four microtubule binding repeats (1N4R) encoding the human P301S mutation under the mouse prion protein promote (The Jackson Laboratory, 008169) were crossed with *Ch25h* ^−/−^ mice (The Jackson Laboratory, 037647) to generate *Ch25h^+/−^ Tau+* mice, and subsequent crossing of F1 litters generated *Ch25h^+/+^ Tau+, Ch25h^−/−^ Tau+,* mice and their corresponding non-*Tau* littermates. Both male and female mice were used for behavioral, histological, and biochemical analyses. Only female mice were use for transcriptomic and lipidomic analyses. Mice underwent behavioral testing at 7–8 months of age and harvested at 9– 10 months of age for pathology and various studies. All experimental procedures involving mice were conducted in accordance with ethical guidelines and were approved by the Institutional Animal Care and Use Committee of Weill Cornell Medicine.

### Human brain samples

Tissues used in this study were Brodmann area 9 (BA9) frontal cortices obtained postmortem from age-matched individuals with Alzheimer’s disease and non-demented controls. All samples were obtained from the Mount Sinai Brain Bank. Brains were donated with consent from the next-of-kin or an individual with the legal authority to provide such permission. The brain tissues used in this study are considered de-identified and therefore do not constitute “human subjects” research under institutional policy, and are not subject to IRB oversight. The Institutional Review Board determined that clinicopathologic studies using de-identified postmortem tissue samples are exempt according to 45 CFR 46.104(d)(2). Additional information about donor characteristics is provided in Supplementary Table 5.

### Mouse and human brain immunohistochemistry and imaging

For mouse brain tissue, Dulbecco’s phosphate-buffered saline (DPBS) was used for immunohistochemistry. Four brain sections per mouse that contain a series of anterior to posterior hippocampus were washed to remove cryoprotectant and then permeabilized by 0.5% Triton X-100. After blocking in 5% normal goat serum (NGS) for 1 h, brain sections were incubated with primary antibodies in the same blocking buffer overnight at 4 °C. Sections were then washed by DPBS containing 0.1% Tween-20 and incubated with Alexa-conjugated secondary antibodies for 1 h at room temperature in blocking buffer. After washing, sections were mounted on glass slides with ProLong Gold Antifade Mounting media.

For human brain samples, slides were placed for 10 min at 60°C in an oven and then de-paraffinized by washing with xylene three times for 5 min, 100% ethanol two times for 2 min, 95% ethanol for 2–5 min, and deionized water 3 times for 2 min. For antigen retrieval, the slides were placed in working Citrate buffer (Electron Microscopy Sciences, Cat # 64142-08) and placed in a pressure cooker (Cuisinart) at high pressure for 15 min. The slides were cooled to RT and washed with cool deionized water three times for 2 min. Sections were washed with 1x PBS three times for 2 min and incubated with 1x TrueBlack (Biotium) in 70% ethanol for 30 s. The reaction was stopped by placing the slides in 1x PBS and further washing with 1x PBS three times for 2 min. Sections were blocked with 5% normal donkey serum in 1x PBS for 1 h and incubated with mouse anti-MC1 Tau (1:500, Peter Davies, Albert Einstein College of Medicine, Cat # MC1), rabbit anti-VGAT (1:600,millipore, ab5062), chicken anti-MAP2 (1:1000,Novus Biological, Cat # NB300-213), rabbit anti-STING (1:500, Cell Signaling Technology, Cat #50494), mouse anti-GM130 (1:600, BD Transduction Laboratories, Cat #610822), rabbit anti-GFAP (1:800, Abcam, Cat #ab7260), goat anti-IBA1 (1:600, Abcam, Cat #ab5076), rat anti-P2RY12 (1:600, Biolegend, Cat #848002), mouse anti-APOE(HJ6.3)(1:500, David Holtzman), rabbit anti-SYNAPTOPHYSIN (D35E4) (1:1,000, Cell Signaling Technology, Cat #5461), mouse anti-PSD95 (1:1,000, Abcam, Cat # ab2723), and rat anti MHC Class II-Alexa Fluor 594 (1:600, Biolegend, Cat #107650). diluted in 1% normal donkey serum in 1x PBS solution overnight in a humidified slide chamber. The following day, the slides were then washed in 1x PBS three times for 2 min and incubated with AF 488 donkey anti-rabbit (1:500) and AF 568 donkey anti-rat (1:500) diluted in 1% normal donkey serum in 1x PBS solution for 1 h in a humidified slide chamber. The slides were washed in 1x PBS three times for 2 min and incubated with Hoescht 33342 diluted (1:1000, Thermo Fisher Scientific; Cat # 62249) in 1x PBS for 15 min. The slides were washed in 1x PBS three times for 2 min and covered with Vectashield mounting medium without DAPI (Vector Laboratories; Cat # H-1000-10). Three imaging fields in the gray matter of the same section were captured using Zeiss Apotome 3 microscope and SLIT2 mean intensity of each section was quantified using analyzed with ImageJ (NIH, RRID:SCR_003070).

The primary antibodies used for immunohistochemistry were as follows(antibody). Images for MC1 and IBA1 quantification were acquired on Zeiss microscope using 20x objective and analyzed with ImageJ (NIH). All images were first set the threshold manually, then the auto-measurements were performed by using the macros program in ImageJ. Regions of interest including the hippocampus and cortex were hand-traced. MC1+ areas were measured by ImageJ, whereas OLIG2+ cell numbers were counted with the Analyze Particles function. 3D structure of microglia was reconstructed using the Imaris 10.0(RRID:SCR_007370) software as described before ^108^. Experimenters performing imaging and quantification were blinded.

### Brain volumetric analysis

Brain volumetric quantification was performed following a previously described method with modifications. Hippocampal sections spanning from bregma ∼1.3 mm to ∼3.1 mm were collected from one series of brain sections (approximately 10–12 sections per animal, spaced 240 mm apart). Sections were mounted and rinsed in Milli-Q water for 1 min, then stained with 0.1% Sudan Black (Abcam, ab146284) in 70% ethanol at room temperature for 30 min. After staining, sections were washed three times in 70% ethanol (10s each), followed by three washes in Milli-Q water (2 min each), and were cover slipped with VectaMount mounting media (Vector laboratories, H-5501). Slides were imaged on a Keyence BZ-X700 microscope using a 34× objective under brightfield with XY stitching. Regions of interest were manually traced in ImageJ, and area measurements were obtained. Volumes were calculated for each animal using the formula: Volume = (sum of area) × 240 µm.

### Western blotting

Total brain cortex lysates were prepared in radioimmunoprecipitation assay buffer (RIPA) [1% NP-40, 0.5% sodium deoxycholate, and 0.1% sodium dodecyl (lauryl) sulfate]. Protein (20 mg) was separated by a 12% SDS-PAGE gel, then transferred to a polyvinylidene difluoride (PVDF) membrane. After blocking in TBS buffer (20 mM Tris-HCl, 150 mM sodium chloride) containing 5% (wt/vol) nonfat dry milk for 1 h at room temperature, the membranes were then probed with proper primary and secondary antibodies, which was followed by developing with Super Signal West Pico chemiluminescent substrate (34577; Thermo Scientific, Rockford, IL). Data analysis was performed by Image lab 6.1 (Bio-Rad, Hercules, CA, RRID:SCR_014210). The following primary antibodies were used: rabbit anti-SYNAPTOPHYSIN (D35E4) (1:1,000, Cell Signaling Technology, Cat #5461), mouse anti-PSD95 (1:1,000, Abcam, Cat # ab2723), mouse anti-total Tau (HT7)(1:3,000, Thermo Fisher Scientific, Cat #MN1000), rabbit anti-STAT1 (1:1,000, Cell Signaling Technology, Cat #14994), rabbit anti-STING (1:1,000, Cell Signaling Technology, Cat #50494), rabbit anti-Phospho-STING (Ser365) (1:1,000, Cell Signaling Technology, Cat #72971), rabbit anti-GAPDH (1:5,000, Cell Signaling Technology, Cat # 2118), and rabbit anti-Beta3-Tubulin (1:5,000, Cell Signaling Technology, Cat # 5568). The following secondary antibodies were used: HRP-goat anti-Mouse IgG (1:2,000, Jackson, Cat # 115-035-146), HRP-goat anti-Rabbit IgG (1:2,000, Jackson, Cat # 111-035-144).

### Mouse Behavioral Testing

The Morris Water Maze test was used to assess spatial memory acquisition and retention using well-defined protocols^109^. Mice were acclimated to the testing environment for at least 1 hour before testing. Animals were randomized by genotype, and the experimenter was blinded to group assignments during testing. In brief, mice were trained to locate a hidden platform (20 cm diameter) in an open circular pool (200 cm diameter) filled with 21 °C water rendered opaque with nontoxic poster paint. A digital camera mounted above the maze recorded the animals’ movements throughout testing. Four trials were conducted per day for 5 consecutive days. Each trial began from a different starting point (N, E, SE, NW), while the platform remained in a fixed position (NE). Each trial lasted 60 s and was followed by a 30 s period during which mice were allowed to remain on the platform to reinforce their memory of its location. After the 5-day acquisition phase and a 24-hour interval, mice underwent a probe (retention) trial in which the platform was removed. The latency to reach the former platform location, the time spent in the target quadrant, and the number of crossings over the previous platform position were analyzed using the Animal Tracker plugin in ImageJ (Version 2.9.0, National Institutes of Health).

### Primary microglial culture

Following established protocol^110^, primary microglia were isolated from the hippocampi and cortices of 0-3-day-old mouse pups. The isolated brain tissues were rinsed with Dulbecco’s Phosphate-Buffered Saline (DPBS), and the meninges were carefully removed. Subsequently, the brain tissues were minced, followed by treatment with 0.05% trypsin at 37 °C for 20 minutes. The trypsinization process was halted by adding 20% FBS/DMEM media, after which the digested tissues were gently triturated to generate a cell suspension. This suspension was then subjected to centrifugation at 200 x g for 15 minutes, and the pellet was resuspended in 10% FBS/DMEM. The resuspended cells were plated onto T-75 flasks coated with poly-D-lysine (PDL), facilitating the formation of mixed glial cultures. These cultures were maintained in 10% FBS/DMEM supplemented with 5ng/ml granulocyte-macrophage colony-stimulating factor (GM-CSF). By the twelfth day, when the cultures had reached confluence, microglia were isolated from the glial layer by subjecting the flasks to gentle shaking at 400 rpm for a duration of 2 hours. The microglia that floated were subsequently seeded onto plates coated with PDL at a density of 75,000 cells/cm^2^. They were then cultured in 10% FBS/DMEM without GM-CSF for a 24-hour period before being employed in assays involving the phagocytosis and processing of tau fibrils.

### Bulk RNA sequencing

Freshly perfused mouse brains were dissected to isolate the cortices. The cortices were flash-frozen and then stored at -80 °C. For RNA extraction, the cortices were thawed on ice for a duration of 30 minutes and then RNA isolation from the cortex tissue was carried out following the manufacturer’s protocol (PureLink™ RNA Mini Kit, Thermo Fisher). The isolated RNA samples were then sent to the Weill Cornell Medicine Genomics Core for assessment of RNA quality and integrity. Following successful quality control, RNA-seq libraries were prepared for sequencing using the NovaSeq platform.

### Isolation of nuclei from frozen mouse brain tissue

The protocol for isolating nuclei from frozen mouse brain tissue was adapted from previous studies with modifications ^111,112^. All procedures were done on ice or at 4 °C. In brief, mouse brain tissue was placed in 1,500 µl of nuclei PURE lysis buffer (Sigma, NUC201-1KT) and homogenized with a Dounce tissue grinder (Sigma, D8938-1SET) with 15 strokes with pestle A and 15 strokes with pestle B. The homogenized tissue was filtered through a 35 µm cell strainer and was centrifuged at 600 × g for 5 min at 4 °C and washed three times with 1 ml of PBS containing 1% BSA, 20 mM DTT, and 0.2 U µl^−1^ recombinant RNase inhibitor. Then the nuclei were centrifuged at 600 × g for 5 min at 4 °C and resuspended in 500 µl of PBS containing 0.04% BSA and 1× DAPI, followed by FACS sorting to remove cell debris. The FACS-sorted suspension of DAPI-stained nuclei was counted and diluted to a concentration of 1,000 nuclei per microliter in PBS containing 0.04% BSA.

### Droplet-based single-nuclei RNA-seq

For droplet-based snRNA-seq, libraries were prepared with Chromium Single Cell 3’ Reagent Kits v3 (10× Genomics, PN-1000075) according to the manufacturer’s protocol. cDNA and library fragment analysis were performed using the Agilent Fragment Analyzer systems. The snRNA-seq libraries were sequenced on the NovaSeq 6000 sequencer (Illumina) with 100 cycles. Gene counts were obtained by aligning reads to the mouse genome (mm10) with Cell Ranger software (v.3.1.0) (10× Genomics). To account for unspliced nuclear transcripts, reads mapping to pre-mRNA were counted. Cell Ranger 3.1.0 default parameters were used to call cell barcodes. We further removed genes expressed in no more than three cells, cells with a unique gene count over 4,000 or less than 300, and cells with a high fraction of mitochondrial reads (>5%). Potential doublet cells were predicted and removed using DoubletFinder ^53^ for each sample.

### Ambient RNA removal using Cellbender^113^

The raw_feature_bc_matrix.h5 file was generated using Cell Ranger software. This file was then processed using the Cellbender remove-background command to create a new matrix.h5 file. Subsequently, this file was read into Seurat, where the raw data underwent further cleaning with DoubletFinder and SoupX^114^. After these steps, the file was utilized in the Seurat pipeline.

### Sn-RNAseq data analysis by Seurat package

Normalization and clustering were done with the Seurat package v4.0.0. In brief, counts for all nuclei were scaled by the total library size multiplied by a scale factor (10,000), and transformed to log space. A set of 2,000 highly variable genes were identified with FindVariableFeatures function based on a variance stabilizing transformation (vst). Principal component analysis (PCA) was done on all genes, and t-SNE was run on the top 15 PCs. Cell clusters were identified with the Seurat functions FindNeighbors (using the top 15 PCs) and FindClusters (resolution = 0.1). For each cluster, we assigned a cell-type label using statistical enrichment for sets of marker genes and manual evaluation of gene expression for small sets of known marker genes. The subset() function from Seurat was used to subset each cell types. Differential gene expression analysis was done using the FindMarkers function and MAST ^115^. For pseudobulk analyses, we aggregated the expression values from all nuclei from the same cell type for genotype dependent differential expression.

### Gene network and functional enrichment analysis

Gene network and functional enrichment analysis were performed by QIAGEN’s Ingenuity® Pathway Analysis (IPA®, QIAGEN Redwood City, www.qiagen.com/ingenuity, Version 01-22-01) or by GSEA with molecular signatures database (MSigDB) ^116,117^. Significant DEGs and their log_2_fold change expression values and FDR were inputted into IPA for identifying canonical pathways, biological functions, and upstream regulators. Significant DEGs were input into GSEA (http://www.gsea-msigdb.org/gsea/msigdb/annotate.jsp) to identify hallmark and gene ontology terms. The p-value, calculated with the Fischer’s exact test with a statistical threshold of 0.05, reflects the likelihood that the association between a set of genes in the dataset and a related biological function is significant. A positive or negative regulation z-score value indicates that a function is predicted to be activated or inhibited.

### iPSC Differentiation and K18 seeding

We used human 4R or 4R-P301S iPSC homozygous lines described in our previous study^56^. Human iPSC lines were maintained in Essential 8 or mTeSR medium and passaged twice before initiating differentiation. For pre-differentiation (day –3 to 0), iPSCs were dissociated with Accutase and seeded on Matrigel-coated plates in pre-differentiation medium: KnockOut DMEM/F12 (Thermo Fisher Scientific; Cat. No. 12660012) supplemented with 1x non-essential amino acids (Thermo Fisher Scientific; Cat. No. 11140050), 1xN2 (Thermo Fisher Scientific; Cat. No. 17502048), 10 ng/mL BDNF (PeproTech; Cat. No. 450–02), 10 ng/mL neurotrophin-3 (NT-3, PeproTech; Cat. No. 450–03), 1 µg/mL laminin (Trevigen; Cat. No. 3446–005-01), 2 µg/mL doxycycline (Sigma-Aldrich; Cat. No. D9891), and 10 µM ROCK inhibitor (Cayman Chemical Cat. No. 10005583). Three days before differentiation, culture media were completely replaced with pre-differentiation medium supplemented with BDNF/NT3/laminin/doxycycline with ROCK inhibitor removed.

On day 0, cells were dissociated with Accutase and replated at 3.8 × 10^5^ /well in poly-D-lysine/laminin-coated 12-well plates (Corning Cat. No. 356470) or 1.5×10^5^/coverslip coated with poly-D-lysine/laminin in 24-well poly-D-lysine precoated tissue culture plates. The cells were maintained in maturation medium (50% Neurobasal-A (Thermo Fisher Scientific; Cat. No. A3582901), 50% DMEM/F12 (Thermo Fisher Scientific; Cat. No. 11320033), 0.5× N2, 0.5× B27 (Thermo Fisher Scientific; Cat. No. A35828–01), NEAA, 0.5xGlutaMAX (Thermo Fisher Scientific; Cat. No. 35050061), supplemented with 10ng/ml BDNF, 10ng/ml NT3, 1ug/ml mouse laminin, and 2ug/ml doxycycline. Cultures were maintained by half-medium replaced every other day using maturation medium containing BDNF and NT3. On day 7, the neurons were treated with or without 1.5ug/ml K18 tau in the presence of 0, 0.1, 0.2 or 2.5uM 25-HC dissolved in ethanol. The neurons were collected for biochemical or fixed using 4% PFA for immunostaining after 14 days treatment.

### DMXAA Preparation and Administration

DMXAA was freshly prepared on the morning of the experiment at a concentration of 3.75 mg/mL in a 5:80:15 (v/v/v) mixture of DMSO, PEG400, and 45% hydroxypropyl-β-cyclodextrin (HPβCD). Mice were intraperitoneally injected with 25 mg/kg DMXAA or an equivalent volume of vehicle control. All injections were performed as an acute 6-hour treatment, and both the injection time and administered volume (calculated according to individual body weights) were recorded. Following 6 hours post-injection, animals were euthanized and tissues were collected for downstream analyses.

### Quantitative real-time PCR of mouse brain tissue

Mouse hippocampal tissue of Ch25h+/+ or-/-treated with vehicle or DMXAA was homogenized, and total RNA was extracted using the PureLink RNA Mini Kit (Invitrogen, Cat. #12183025), followed by genomic DNA removal with the RNase-Free DNase Set (Qiagen, Cat. #79254). After cDNA synthesis, RNA expression levels of Irf3 (Cat. #4331182; Assay ID: Mm00516784_m1), Irf7 (Cat. #4331182; Assay ID: Mm00516793_g1), and C1qb (Cat. #4331182; Assay ID: Mm01179619_m1) were quantified using TaqMan assays, that of Stat1 was validated in the presence of SsoAdvanced Universal SYBR Green Supermix (Biorad, 1725271) on a Bio-Rad CFX96 Touch Real-Time PCR system. The relative expression of the target genes was normalized to GAPDH mRNA. Stat1 primer sequences: Forward: 5’-TCACAGTGGTTCGAGCTTCAG-3’; Reverse: 5’-CGAGACATCATAGGCAGCGTG-3’.

### LC-MS/MS lipidomic analysis of mouse brain tissues

#### For the lipid profiling of *Tau-* and *Tau+* tissue

Lipidomic analyses were performed by the Biomarker Core at Columbia as previously described using Ultra Performance Liquid Chromatography-Tandem Mass Spectrometry (UPLC-MSMS) after extraction of lipids using a modified Bligh and Dyer method. Lipid extract was spiked with appropriate internal standards and analyzed on the Agilent 1260 Infinity HPLC integrated to Agilent 6490A triple-quadropole mass spectrometer controlled by Masshunter v 7.0 (Agilent Technologies, Santa Clara, CA) as previously described^90,118,119^. To quantify the relative molar amounts of the lipid species, multiple reaction monitoring (MRM) transitions with both positive and negative ionization modes, referencing internal standards (Avanti Polar Lipids, Alabaster, AL). The total molar amount of all lipid species was summed from each spectra and used to compare the relative molar percent for each lipid. The data are presented as relative mean molar percent with standard error (mean ± S.E).

#### For the lipid profiling of cortical tissues from a mouse cohort of four genotypes (*Ch25h+/+, Ch25h-/-, Ch25h+/+ Tau+, Ch25h-/-Tau+*) and from the postmortem human brains

weighed tissue was homogenized in with ice-cold phosphatebuffered saline using a bead mill homogenizer. Tissue lysates (∼50 μg) were transferred to Pyrex glass tubes with a PTFE-liner cap. Lipids were then extracted by the Folch method (reff). Briefly, 6 mL of ice-cold chloroform, methanol (2:1 v/v), and 1.5 mL of water were added to the samples and tubes were vortexed thoroughly to mix the samples homogenously with a polar and non-polar solvent. SPLASH LIPIDOMIX internal standards (Avanti Polar Lipids) were spiked in before the extraction. The organic phase of each sample was normalized by internal standards. After vortexing, samples were centrifuged for 20 min at 1000 rpm at 4°C to separate the organic and inorganic phases. Using a sterile glass pipette, the lower organic phase was transferred into a new glass tube, taking care to avoid the intermediate layer of cellular debris and precipitated proteins. The samples were dried under nitrogen flow until the solvents were completely dried. Samples were resuspended in 150 μL of Isopropanol: Acetonitrile: water (60:35:5) and stored in -80°C until mass spectrometer (MS) analysis. For Ganglioside extraction. ∼100 μg homogenized tissues were extracted with methanol and spun down at 1000 x g for 20 mins to pellet proteins. Solvent layer containing gangliosides and other metabolites was collected in a fresh glass tube and dried under N2 stream. Dried extract was reconstituted in 1 mL LC-MS grade water and desalted by SOLA HRP SPE 30mg/2mL 96 well plate 1EA (Thermo Scientific #60509-001). Desalting cartridges were cleaned 3 times with 1mL methanol and equilibrated 3 times with LC-MS grade water. Next, the extracts dissolved in water were loaded onto the cartridge, washed with 2mL of water then gangliosides were eluted with 3 mL methanol. Eluate was dried under nitrogen flow and reconstituted in 300:150:50 of MeOH: water: chloroform.

Lipids were separated using ultra-high-performance liquid chromatography (UHPLC) coupled with tandem mass spectrometry (MS/MS). UHPLC analysis was conducted on a C30 reverse-phase column (Thermo Acclaim C30, 2.1 x 150 mm, 2.6 μm) maintained at 50°C and connected to a Vanquish Horizon UHPLC system, along with an OE240 Exactive Orbitrap MS (Thermo Fisher Scientific) equipped with a heated electrospray ionization probe. Each sample (2 μL) was analyzed in both positive and negative ionization modes. The mobile phase included 60:40 water:acetonitrile with 10 mM ammonium formate and 0.1% formic acid, while mobile phase B consisted of 90:10 isopropanol:acetonitrile with the same additives. The chromatographic gradient involved: Initial isocratic elution at 30% B from -3 to 0 minutes, followed by a linear increase to 43% B (0-2 minutes), then 55% B (2-2.1 minutes), 65% B (2.1-12 minutes), 85% B (12-18 minutes), and 100% B (18-20 minutes). Holding at 100% B from 20-25 minutes, a linear decrease to 30% B by 25.1 minutes, and holding from 25.1-28 minutes. Flow rate of 0.26 mL/minute, injection volume of 2 μL, and column temperature of 55°C. Mass spectrometer settings included an ion transfer tube temperature of 300°C, vaporizer temperature of 275°C, Orbitrap resolution of 120,000 for MS1 and 30,000 for MS2, RF lens at 70%, with a maximum injection time of 50 ms for MS1 and 54 ms for MS2. Positive and negative ion voltages were set at 3250 V and 2500 V, respectively. Gas flow rates included auxiliary gas at 10 units, sheath gas at 40 units, and sweep gas at 1 unit. High-energy collision dissociation (HCD) fragmentation was stepped at 15%, 25%, and 35%, and data-dependent tandem MS (ddMS2) ran with a cycle time of 1.5 s, an isolation window of 1 m/z, an intensity threshold of 1.0e4, and a dynamic exclusion time of 2.5 s. Full-scan mode with ddMS2 was performed over an m/z range of 250-1700, with EASYICTM used for internal calibration. The raw data were processed and aligned with LipidSearch 5.1, using a precursor tolerance of 5 ppm and a product tolerance of 8 ppm. Further filtering and normalization were conducted using an in-house app, Lipidcruncher^120^. Semi-targeted quantification was performed by normalizing the area under the curve (AUC) to the AUC of internal standards and further normalized with the total quantified protein.

### LC-MS/MS lipidomic analysis of human iPSC-derived neurons

Lipids were extracted from samples of human iPSC-derived neurons using isopropanol according to a published method^121^. The extracts were clarified by centrifugation and dried down using a SpeedVac. The dried sample was reconstituted using acetonitrile/isopropanol/water 65:30:5 prior to LC-MS analysis. Chromatographic separation was performed on a Vanquish UHPLC system with a Cadenza CD-C18 3 µm packing column (Imtakt, 2.1 mm id x 150 mm) coupled to an Orbitrap Exploris 240 mass spectrometer (Thermo Scientific) via an Ion Max ion source with a HESI probe (Thermo Scientific). The mobile phase consisted of buffer A: 60% acetonitrile, 40% water, 10 mM ammonium formate with 0.1% formic acid and buffer B: 90% isopropanol, 10% acetonitrile, 10 mM ammonium formate with 0.1% formic acid. The LC gradient was as follows: 0–1.5 min, 32% buffer B; 1.5-4 min, 32-45% buffer B; 4-5 min, 45-52% buffer B; 5-8 min, 52-58% buffer B; 8-11 min, 58-66% buffer B; 11-14 min, 66-70% buffer B; 14-18 min, 70-75% buffer B; 21-25 min, isocratic 97% buffer B, 25-25.1 min 97-32% buffer B; followed by 5 min of re-equilibration of the column before the next run. The flow rate was 200 μL/min. A data-dependent mass spectrometric acquisition method was used for lipid identification. In this method, each MS survey scan was followed by MS/MS scans performed on the most abundant ions. Data was acquired in positive and negative modes. Parameters for MS scans: resolution, 60,000; automatic gain control target, 1e6; maximum injection time, 200 ms; scan range, 250-1800 m/z. Parameters for MS/MS scans: resolution, 15,000; automatic gain control target, standard; maximum injection time, 50 ms; isolation window, 1.5 m/z; NCE, stepped 20,30 and 40. The LC-MS files were processed using MS-DIAL software (version 4.9) for lipid identification and relative quantitation after normalized by total proteins.

### TUNEL assay

Human neurons under different treatment conditions for 14 days were fixed with 4% paraformaldehyde in PBS for 1 h at room temperature, permeabilized with 0.1% Triton X-100 in 0.1% sodium citrate for 2 min on ice and then incubated with the TUNEL reaction mixture from the In Situ Cell Death Detection Kit, TMR Red (Sigma-Aldrich) for 60 min at 37 °C in a humidified dark chamber. After washing with PBS, samples were mounted in VECTASHIELD mounting medium with DAPI (Vector Laboratories; Cat # H-1200-10) and examined under a fluorescence microscope (excitation 520–560 nm; emission 570–620 nm). The TUNEL signals were quantified by ImageJ (NIH, RRID:SCR_003070).

## Statistical Analyses

The sample size for each experiment was determined based on previous publications^122,123^. All in vitro experiments were performed with a minimum of three biological replicates. Mean values from at least three independent experiments were used for computing statistical differences. All *in vivo* experiments were performed with a minimum of four mice (for snRNAseq) or 6-12 mice/genotype. In vivo data were averaged using the appropriate unit: per mouse for IHC staining. and mean values were used for computing statistical differences. Data visualization was done with Graphpad and R package ggplot2. Statistical analyses were performed with Graphpad prism 10 (t-test, one-way and two-way ANOVA) (Graphpad, San Diego, California). Values are reported as mean ± standard error of the mean (SEM). Mann–Whitney test was used when the normality test is not passed. One-way ANOVA was used to compare data with more than two groups. Two-way ANOVA was used for groups with different genotypes and/or time as factors. Tukey’s and Sidak’s post-test multiple comparisons were used to compare the statistical difference between designated groups. All P-values of enrichment analysis are calculated by right-tailed Fisher’s exact test. P < 0.05 was considered statistically significant.

## Acknowledgments

This work was supported by the NIH (1R01AG096902-01 to W.L., L.G. and S.G.; R01AG072758 (to L. G.), R01AG079557-01 (to L. G.),1R01AG079291-01A1 (to L.G.), and R01AG074541( to L.G.), the Rainwater Charitable Foundation ( to L. G.), Freedom Together Foundation (to L.G.), Cure Alzheimer’s Fund (to L.G.), and Hevolution Foundation partnership grant (to L.G.). NIH 1R56AG062271-01A1 (to L.B.M.) and National Center for Advancing Translational Sciences, NIH UL1TR001873 (to L.B.M.). Canadian Institutes of Health Research (PJT-190004, to S.A.M.) We acknowledge the help from Dr. Guoan Zhang and Mengmeng Zhu in the Proteomics and Metabolomics Core Facility in Weill Cornell Medicine for lipidomics. We acknowledge Dr. Renu Nandakumar the Columbia University Medical Center Biomarker Core. We acknowledge the Mount Sinai Brain Bank for providing postmortem human brains for this analysis.

## Author contributions

W.L. and L.G conceived the project. W.L., L.G., and H.C. designed experiments. H.C., L.F., Y. A., M.Y.W., J.Z., N.F., P. Y., K.N., Y.S.W., D.Z., T. P., L.B.M. performed experiments or analyses. S.A.M., T.W., and R.F. developed experimental protocols, tools, or reagents. M.Y.W., K.N., and P.Y. maintained the mouse colony. W.L., L.G. and H.C. wrote the manuscript. All authors read and approved the paper.

## Resource availability Lead contact

Further information and requests for resources and reagents should be directed to and will be fulfilled by the lead contact, Wenjie Luo; Li Gan.

## Materials availability

Mouse lines generated in this study are available upon request without restrictions with a completed materials transfer agreement for not-for-profit organizations but may require payment if there is potential for commercial application.

## Data and code availability

Bulk and single-cell RNA sequencing data have been deposited at GEO and will be publicly available as of the date of publication. Accession number for Bulkseq is GSE313022, for single nuclei seq is GSE313021.

## AI Statement

This manuscript was edited by ChatGPT for typographical and grammatical accuracy.

## Supplementary Figures

**Supplementary Figure 1.**
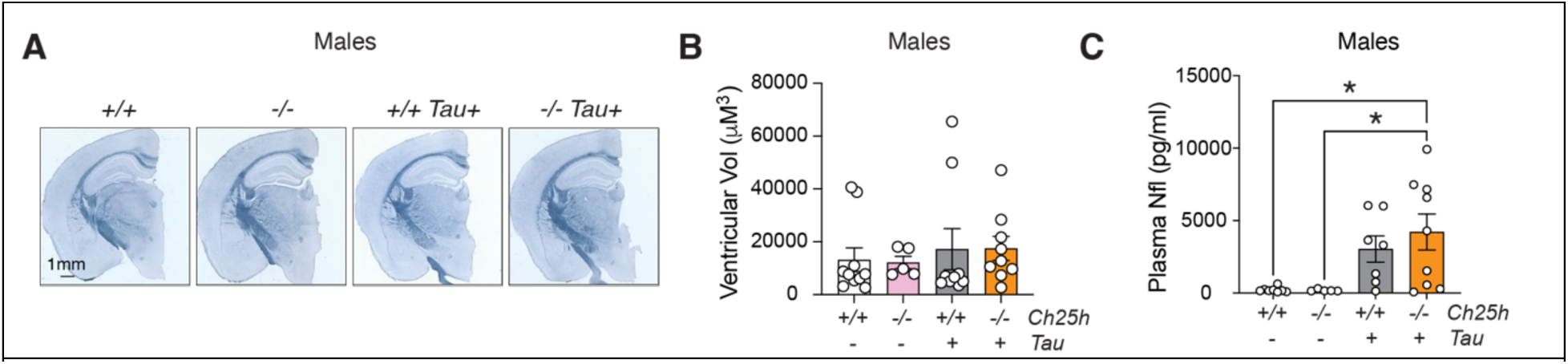
*Ch25h* deficiency does not mitigates neurodegeneration in male tauopathy mice (related to Figure 1) A) Representative Sudan black images of brains from 9- to 10-month-old male mice across four genotypes. B) Quantification of ventricular volume of each mouse brain across four genotypes. C) Quantification of plasma Nfl levels measured by the ELISA assay across four genotypes. Statistical analyses were performed by Two-Way ANOVA followed with Tukey post-hoc test. *p<0.05.

**Supplementary Figure 2.**
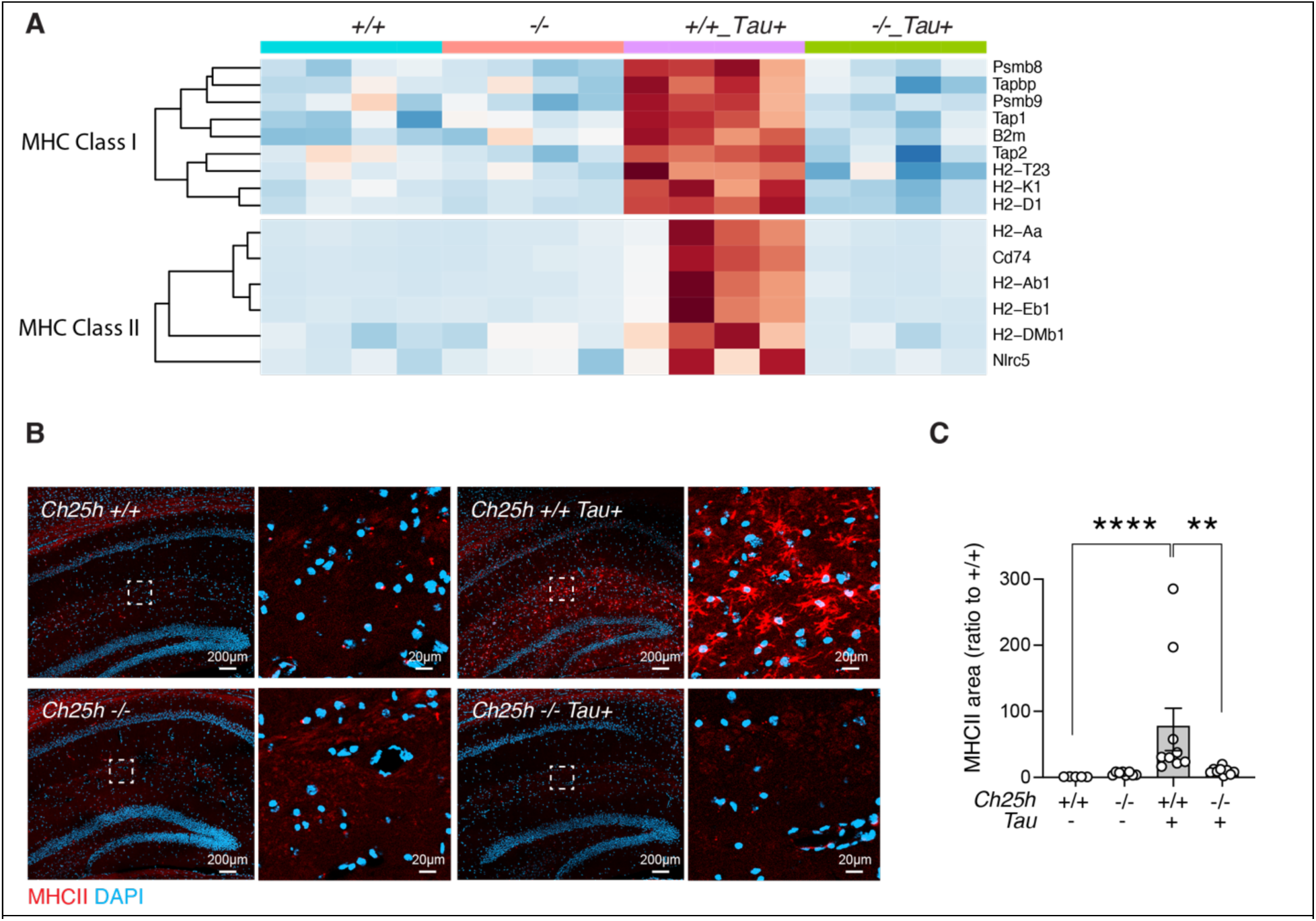
*Ch25h* deletion reduces tau-induced MHC expression in female tauopathy mouse brains (related to. **Figure 2).** A) Heatmap of MHC I and MHC II associated gene expressions in mice across four genotypes. B-C) Representative images (B) and quantification (C) of the MHCII in the hippocampus of mice across four genotypes. n = 5 mice for *Ch25h+/+*, n = 8 mice for *Ch25H -/-,* n = 9 mice for *Ch25h+/+ Tau+,* n = 9 mice for *Ch25h-/- Tau+.* **p<0.01, ****p<0.0001. Two-Way ANOVA followed with Tukey test.

**Supplementary Figure 3.**
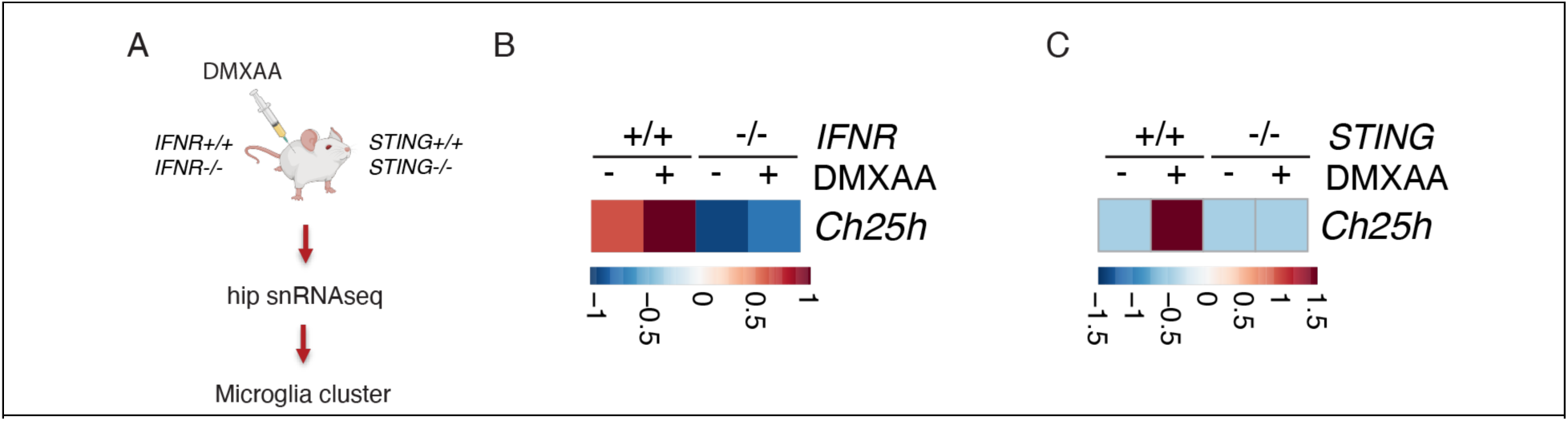
STING–IFN signaling induces microglial *Ch25h* expression (related to. **Figure 3).** A) Schematic of experimental workflow showing 6h post 25mg/kg DMXAA i.p injection followed by single-nucleus RNA sequencing to analyze microglial clusters. B) Heatmap showing *Ch25h* expression in microglia from *IFNR* +/+ and *IFNR* -/- mice treated with or without DMXAA. C) Heatmap showing *Ch25h* expression in microglia from *STING* +/+ and *STING* -/- mice treated with or without DMXAA.

**Supplementary Figure 4:**
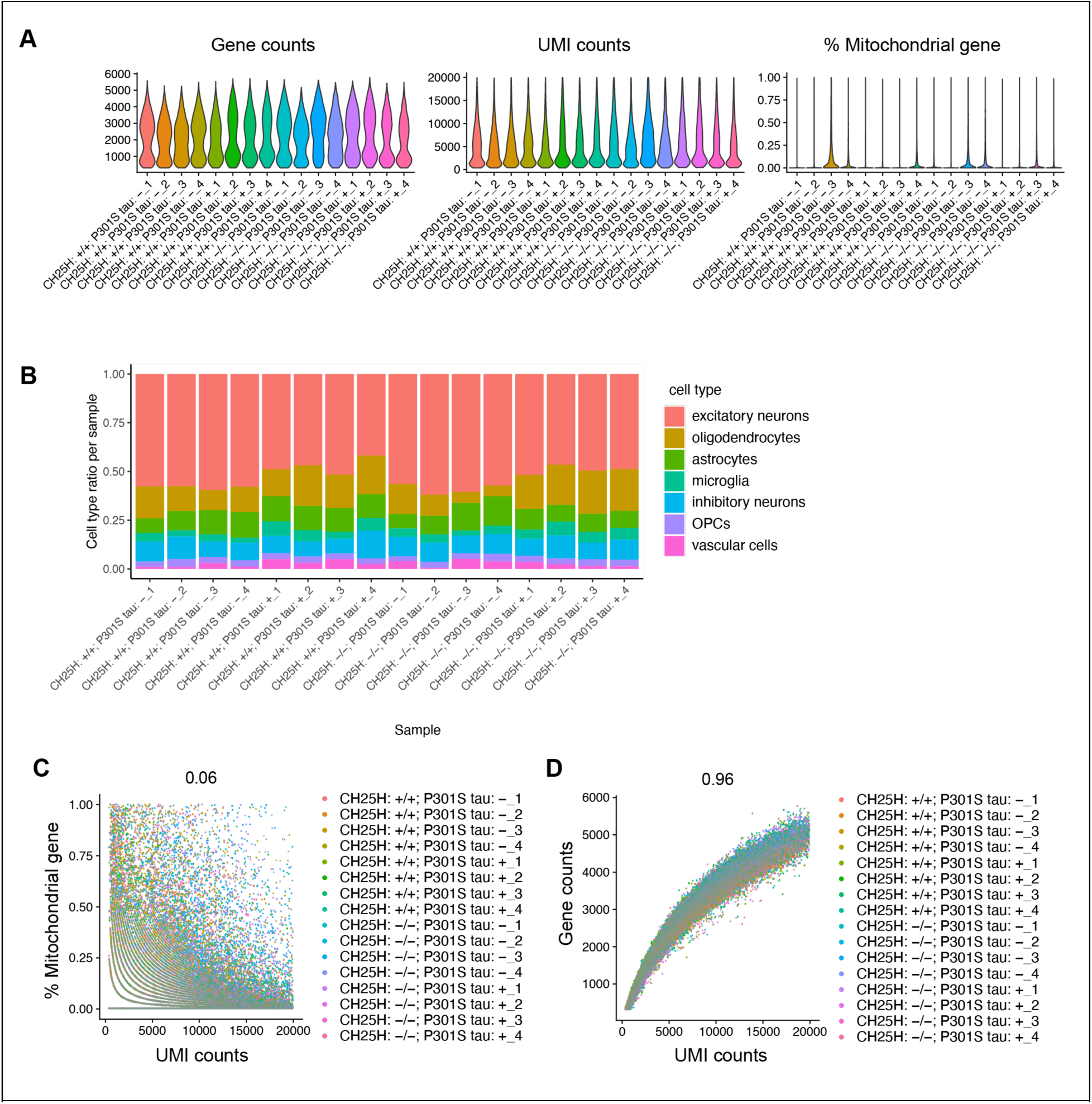
Quality control assessment of snRNAseq of mouse hippocampus across four genotypes (related to. **Figure 4).** A) Quality control plots showing total number of genes, total number of molecules, and equivalent amounts of percent mitochondrial RNA in nuclei used for downstream analyses. B) Cell ratio across all samples. C-D) Correlation between UMI counts and percentage of mitochondrial genes per nuclei (C) and total genes detected (D) for all samples.

**Supplementary Figure 5.**
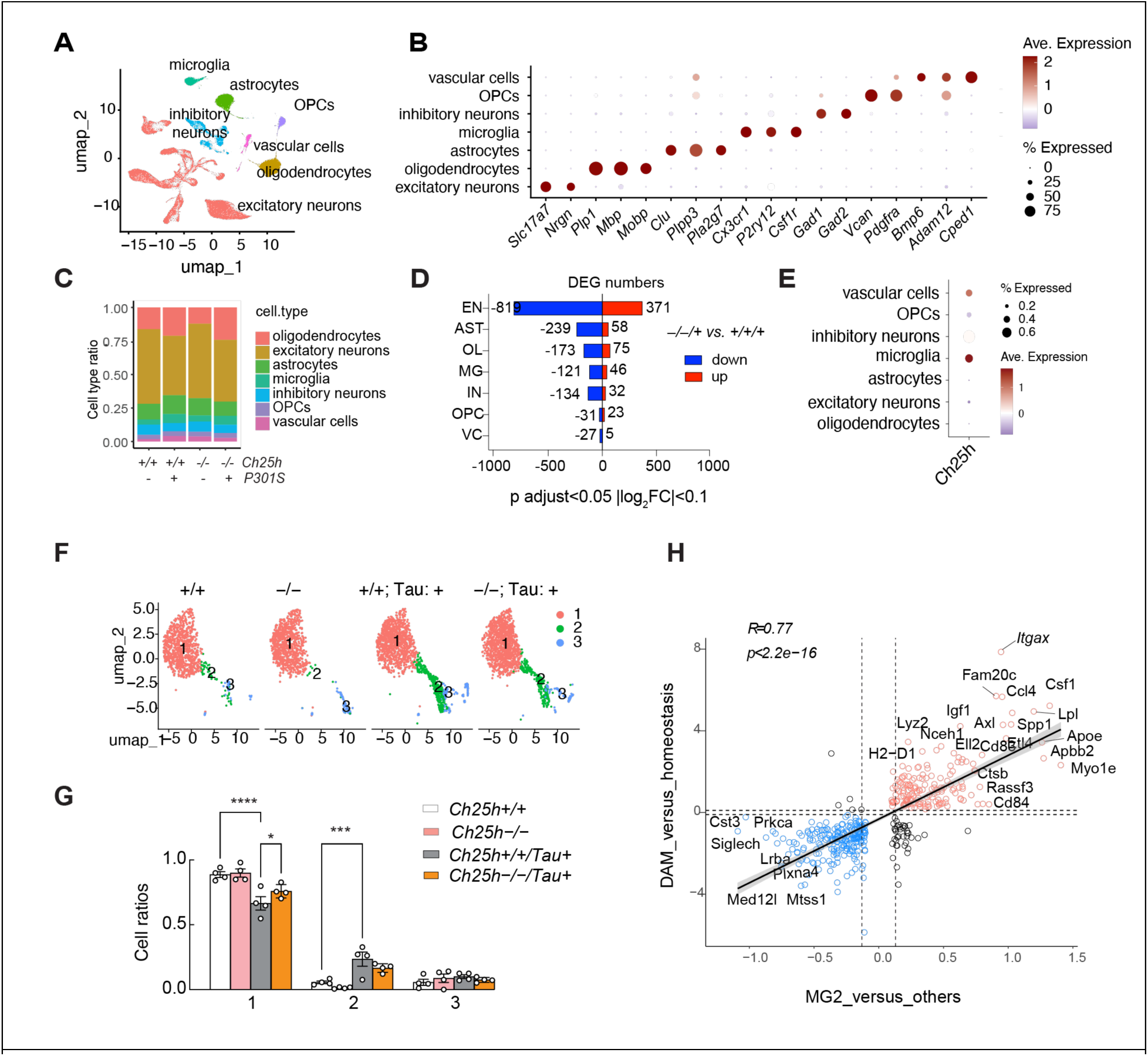
snRNAseq analysis of hippocampus of female tauopathy mice across genotypes (related to. **Figure 4).** A) UMAP dimensional plot showing nuclei colored according to transcriptionally distinct cell clusters identified using Seurat package. B) Summary of marker genes used for cluster classification into different cell types. C) Ratios of different cell types in each genotype. D) The numbers of DEGs in each cell type altered by *Ch25h* deletion in *Tau+* mice. E) Ch25h expression across different cell types. F) UMAP of microglial subclusters across four genotypes. G) Cell ratios of each microglial subclusters across four genotypes. Two-Way ANOVA followed by post-hoc tukey analysis, *p<0.05, ***p<0.001, ****p<0.0001. H) Correlation analysis of MG2 vs other subcluster and DAM vs homeostatic genes.

**Supplementary Figure 6.**
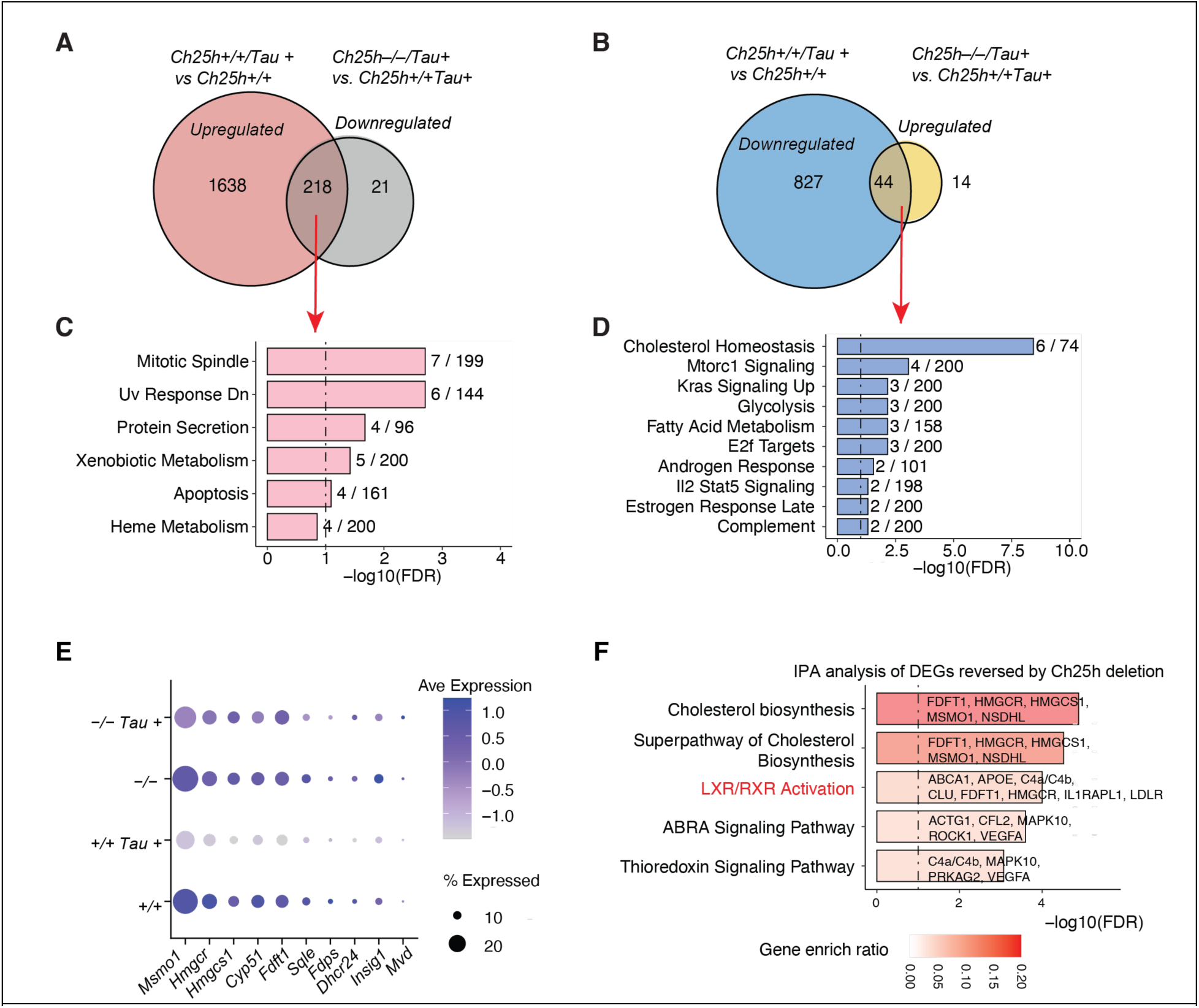
Effects of Ch25h deletion on astrocyte transcriptome in tauopathy mouse brain (related to. **Figure 4)** A-B) Venn diagram identifying changes of astrocytic genes altered by tau and reversed by *Ch25h* deletion in tauopathy mouse brains. C) Top GSEA pathways enriched among DEGs in A (genes upregulated by tau but suppressed by *Ch25h* deletion). D) Top GSEA pathways enriched among DEGs in B (genes downregulated by tau but normalized by *Ch25h* deletion). E) Dot plot showing expressions of genes associated with cholesterol metabolism. G) IPA analysis of DEGs reversed by *Ch25h* deletion in *Tau+* mice.

**Supplementary Figure 7.**
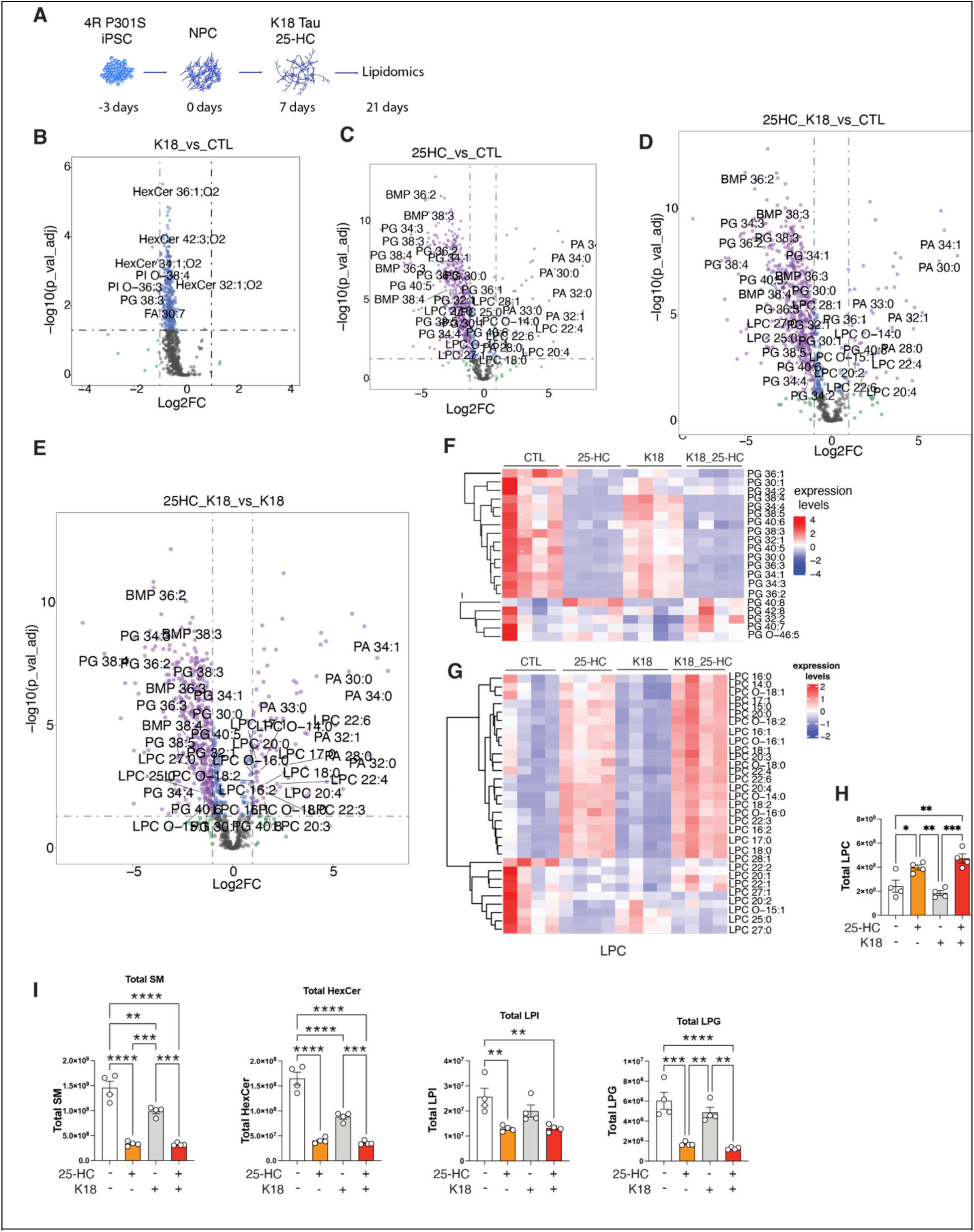
Lipidomic analysis of human neurons (related to. **Figure 6).** A) Schematic of the experimental workflow showing differentiation of 4R P301S iPSC-derived neurons, treatment with tau K18 seeds with or without 2.5 μM 25-HC, and lipidomic profiling at Day 21. B-E) Volcano plots showing lipid species significantly altered by tau seeds, 25-HC treatment, or the combination of tau seeds and 25-HC in human neurons. F-G) Heatmap of phosphatidylglycerol (PG) (F) and lysophosphatidylcholine (LPC) (G) species levels across treatment groups (Vehicle, 25-HC, K18, K18 + 25-HC). H) Quantification of total lysophosphatidylcholine (LPC) levels and heatmap of individual LPC species levels among treatment groups I) Quantification of total sphingomyelin (SM), hexosylceramide (HexCer), lysophosphatidylinositol (LPI), and lysophosphatidylglycerol (LPG) levels in human neurons across all experimental conditions. Statistical analyses in H and I were performed by Two-Way ANOVA followed by tukey posthoc test. *p<0.05, **p<0.01, ***p<0.001, ****p<0.0001.

